# A SI3-σ arch stabilizes cyanobacteria transcription initiation complex

**DOI:** 10.1101/2022.10.06.511230

**Authors:** Liqiang Shen, Giorgio Lai, Linlin You, Jing Shi, Xiaoxian Wu, Maria Puiu, Zhanxi Gu, Yu Feng, Yulia Yuzenkova, Yu Zhang

**Affiliations:** Key Laboratory of Synthetic Biology, CAS Center for Excellence in Molecular Plant Sciences, Shanghai Institute of Plant Physiology and Ecology, Chinese Academy of Sciences, Shanghai 200032, China; University of Chinese Academy of Sciences, Beijing 100049, China; Centre for Bacterial Cell Biology, Biosciences Institute, Faculty of Medical Sciences, Newcastle University, Newcastle upon Tyne NE2 4AX, UK; Department of Biophysics, and Department of Infectious Disease of Sir Run Run Shaw Hospital, Zhejiang University School of Medicine, Hangzhou 310058, China

## Abstract

Multi-subunit RNA polymerases (RNAPs) associate with initiation factors (σ in bacteria) to start transcription. The σ factors are responsible for recognizing and unwinding promoter DNA in all bacterial RNAPs. Here, we report two cryo-EM structures of cyanobacterial transcription initiation complexes at near-atomic resolutions. The structures show that cyanobacterial RNAP forms an ‘SI3-σ’ arch interaction between domain 2 of σA (σ_2_) and the sequence insertion 3 (SI3) in the mobile catalytic domain Trigger Loop (TL). The ‘SI3-σ’ arch facilitates transcription initiation from promoters of different classes through sealing the main cleft and thereby stabilizing RNAP-promoter DNA open complex. Disruption of the ‘SI3-σ’ arch disturbs cyanobacteria growth and stress response. Our study reports the structure of cyanobacterial RNAP and unique mechanism for its transcription initiation. Our data suggest functional plasticity of SI3 and provide foundation for further research into cyanobacteria and chloroplasts transcription.

## INTRODUCTION

Most bacterial RNA polymerase core enzymes are composed of five subunits, including two identical α subunit, one β subunit, one β′ subunit, and one ω subunit (1). Certain Gram-positive bacteria can also contain additional accessory δ and ε subunits (2). Bacterial RNAP core enzyme associates with σ factors to form the RNAP-σ holoenzymes that are capable of initiating gene transcription (3). The domain 2 (σ_2_) and domain 4 (σ_4_) of σ^70^-type σ factors are anchored on the surface of RNAP by the β′ coiled-coil motif and the β flap-tip helix respectively, while the domain 3.2 (σ_3.2_) is thread through the active-site cleft to connect σ_2_ and σ_4_(4−6). In the RNAP holoenzymes, the two structure modules—β′ clamp/σ_2_ and β protrusion/lobe—function as two pincers to guide, load, and restrain DNA in the main left of RNAP. During transcription initiation, σ_2_ nucleates the unwinding process of promoter DNA at the −10 element and recognizes the nucleotide sequence of the unwound −10 element through capturing of the −11 and −7 nucleotides of the non-template strand into respective protein pockets (7-9).

The composition, working mechanism, and basic transcription factors of bacterial RNAP are conserved across most bacterial species, however, the transcription apparatus is unique in cyanobacteria, an ancient, large, and diverse bacteria phylum. Cyanobacterial RNAP has two distinctive features compared with other bacterial RNAPs. First, in contrast to a typical bacterial RNAP containing five subunits (2αββ′ω), the *rpoC* gene encoding the full-length RNAP-β′ subunit is split into two genes in cyanobacteria—*rpoC1* and *rpoC2*, resulting in a six-subunit cyanobacterial RNAP (2αβγβ′ω) (1σ, 11). The *rpoC1* gene encodes the N-terminal half of the full-length RNAP-β′ subunit (*γ* subunit; referred as β′1 subunit) and the *rpoC2* gene encodes the C-terminal half of the full-length β′ subunit (β′ subunit; referred as β′2) (Figs. 1A and S1)(11). Second, cyanobacterial RNAP contains by far the largest SI3 insertion in its β′2 subunit (630 residues in *Synechocystis* sp. PCC 6803; shortened as *Syn*6803 herein) out of all bacterial RNAPs (10, 12). The 630 residues are inserted between the two helices of the TL, a key mobile element in catalytic center of RNAP that is essential for catalysis of phosphodiester-bond formation, NTP discrimination, pausing, and cleavage of backtracked RNA (13-18). TL oscillates between an unfolded (inactive) loop and a folded Trigger Helix (TH) (catalytically active) conformations. *E. coli* RNAP SI3 (188 residues) has been demonstrated to affect multiple events in transcription initiation, transcription pausing, and intrinsic termination through regulating the folding of the TL (19-23). Compared with *E. coli* RNAP SI3, which comprises two sandwich-barrel-hybrid motifs (SBHM), the cyanobacterial RNAP-SI3 domain comprises nine SBHM (12), only one of which shares sequence similarity with *E. coli* RNAP SI3. Notably, chloroplast-encoded RNAP (PEP) retains SI3, alga PEP has a similar-size SI3 and PEP of higher plants is ∼160 amino acid residues larger. To date, only the SI3 structure of *E. coli* RNAP has been reported. The structure and function of the largest SI3 insertion of cyanobacterial RNAP remain unknown.

**Figure 1.**
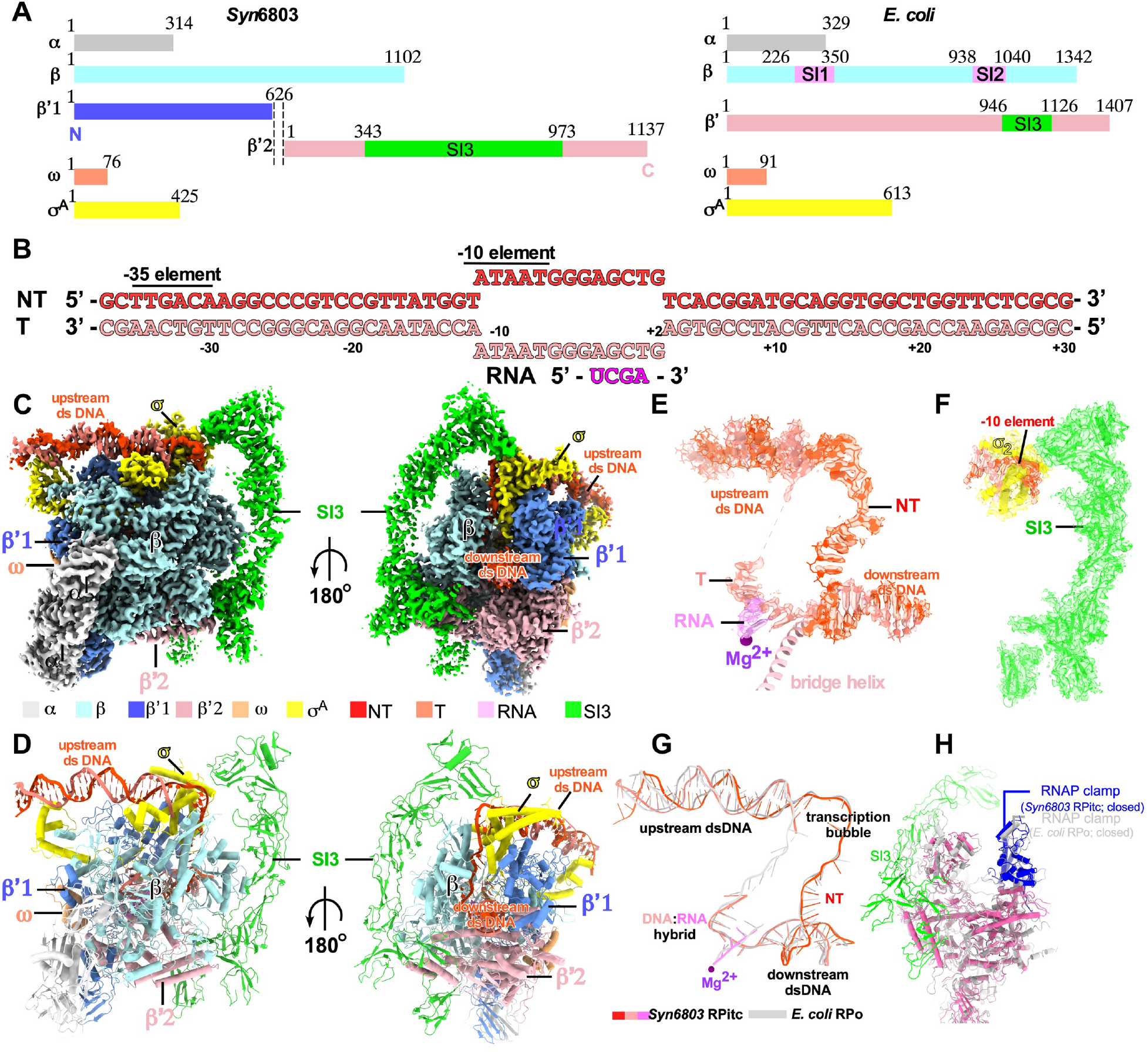
The cryo-EM structure of *Syn*6803 RPitc. **(A)** RNAP subunits of cyanobacteria *Synechocystis* sp. PCC 6803 (left) and *E. coli* (right). **(B)** The nucleic-acid scaffold used in the cryo-EM structure determination. **(C)** Front and back view orientations of *Syn*6803 RPitc cryo-EM map. **(D)** The front and back view orientations of *Syn*6803 RPitc structure. **(E)** The cryo-EM map and model of the nucleic-acid scaffold of *Syn*6803 RPitc. **(F)** the cryo-EM map and model of SI3, σ_2_, and non-template −10 element DNA of *Syn*6803 RPitc. **(G)** Superimposition of the nucleic-acid scaffold between *Syn*6803 RPitc and *E. coli* RPo. **(H)** Superimposition of the clamp domain between *Syn*6803 RPitc and *E. coli* RPo. RNAP subunits and nucleic-acid chains are colored as in the color scheme.

Uniquely for free-living bacteria with complex metabolism, cyanobacteria have reduced repertoire of basic transcription factors. Firstly, cyanobacterial genomes do not encode any secondary channel-binding factors (10), most of which play crucial roles in *E. coli*: for example, the transcription initiation factor, DksA together with (p)ppGpp, represses ribosomal RNA transcription during nutrient-limitation stress (22, 24); and the transcription elongation factors GreA/B facilitate cleavage of RNA in backtracked elongation to restart stalled RNAPs and thus prevent gene expression traffic jams and detrimental collisions with the replication fork (25). We have shown that the intrinsic high hydrolytic activity of cyanobacterial RNA polymerase compensates to a large extent for the absence of these transcription proofreading factors in a recent study (26). Secondly, cyanobacterial genomes don’t encode the termination factor Rho, and thereby are proposed to primarily terminate transcription through the intrinsic termination mechanism. Rho terminates normal and pervasive transcription in *E. coli* (27, 28), and removal of Rho results in severe growth defect in various bacteria (29, 30). It is unknown how cyanobacteria efficiently regulate the transcription termination events without Rho.

To understand the molecular mechanism of gene transcription regulation in cyanobacteria, we took the first step and explored structural basis of the unique transcription apparatus of cyanobacterial RNAP. We determined a cryo-EM structure of *Synechocystis* sp. PCC 6803 transcription initiation complex (*Syn*6803 RPitc) at a resolution of 3.1 Å and a CTP-bound RPitc at a resolution of 3.0 Å. The structures show that the large SI3 domain extends from the bottom of secondary channel to the top of main cleft of RNAP and makes extensive interaction with the rim helices and lobe domain of the RNAP core enzyme and σ factor. Notably, the SI3-head module forms a SI3-σ arch that seals the main cleft of RNAP and stabilizes transcription initiation complexes. Biochemical and genetic evidence suggests importance in RNAP-promoter open complex (RPo) formation of the SI3-σ arch interaction. Our study provides a structural basis for understanding intrinsic properties of cyanobacterial RNAP and a foundation for further exploration of gene transcription regulation in cyanobacteria and chloroplasts.

## RESULTS

### The cryo-EM structure of cyanobacterial RPitc

To obtain a recombinant *Syn*6803 RNAP, we initially co-expressed the six *Syn*6803 RNAP subunits (2α, β, β′1, β′2, and ω subunits) in *E. coli* cells, but failed in obtaining sufficient amounts of recombinant RNAP core enzyme due to poor solubility of the three largest subunits (RNAP-β, β′1, and β′2 subunits). We suspect that the split *Syn*6803 RNAP β′1 and β′2 subunits might be difficult in assembling with other recombinant subunits in *E. coli* cells, therefore we connected *Syn*6803 RNAP-β′1 and -β′2 subunits with a 6-residue flexible linker and were able to obtain functional recombinant *Syn*6803 RNAP in *E. coli* cells for cryo-EM study (Fig. S2A).

The *Syn*6803 RPitc complex was reconstituted using the recombinant *Syn*6803 RNAP, *Syn*6803 σ^A^, a nucleic-acid scaffold composed of a 26-bp upstream dsDNA, a pre-melted 13-bp transcription bubble, a 28-bp downstream dsDNA, and a 4-nt RNA primer (Figs. 1A, 1B, S2B). The structure of *Syn*6803 Rpitc was determined at a resolution of 3.1 Å through a cryo-EM single-particle method (Fig. S3). The cryo-EM map exhibits clear signals for all subunits of RNAP and four major domains (σ_2_, σ_3.1_, σ_3.2_, and σ_4_) of σ^A^ (Figs. 1C and 1D). The cryo-EM map also reveals clear and sharp signals for most nucleotides of upstream (−37 to −13) and downstream (+3 to +30) dsDNA, all nucleotides (−11 to +2) of single-stranded non-template DNA of the transcription bubble, eight nucleotides (−6 to +2) of the single-stranded template DNA of the transcription bubble, and a 4-nt RNA primer base-paired with template DNA in a post-translocated state (Fig. 1E). Our structure shows that in cyanobacterial RPitc, RNAP-σ^A^ holoenzyme adopts a closed conformation, induces near 90° bend of promoter DNA at both junctions of the transcription bubble (Figs. 1G and 1H), and accommodates promoter DNA as other bacterial RNAP-σ^A^ holoenzymes do (Fig. 1G) (9).

Our structure shows that the RNAP-β′1 and RNAP-β′2 subunits are split at a loop region located at the surface of RNAP (Figs. S2D-G). Both split ends of RNAP-β′ subunit (the C terminus of RNAP-β′1 subunit and N terminus of RNAP-β′2 subunit) are well resolved in the cryo-EM map, while the extraneous 6-residue linker is disordered, suggesting that linker doesn’t perturb local structure folds (Fig. S2F). Structure superimposition of *Syn*6803 RNAP with other bacterial RNAP reveals essentially the same structural fold and conformation of the two helices at the split ends (Fig. S2G). Protein sequence alignment suggests that very few residues are deleted or inserted at the C terminus and N terminus of RNAP-β′1 and -β′2 subunits, respectively in various cyanobacteria, even though respective genes encoding the two subunits are separated by ∼0.5 Mbp in certain cyanobacteria (Fig. S1). Altogether, our structure shows that the two largest subunits (β′1 and β′2 subunits) of *Syn*6803 RNAP exhibit structural fold and interaction with the rest of RNAP at the split point similar to that of the unsplit RNAP-β′ subunit.

### Cyanobacterial RNAP-SI3 encloses a large RNAP surface

The cryo-EM map reveals strong signal for *Syn*6803 RNAP-SI3 domain, but local resolution of most regions spans from 4.5 to 7.0 Å except for the sub-regions making contacts with the rest of RNAP holoenzyme (Figs. 1F and S3F). Since the low resolution of RNAP-SI3 domain doesn’t permit *ab initio* model building, we determined a 1.6 Å crystal structure of *Synechococcus elongatus* PCC 7942 (*Syn*7942) RNAP SI3-tail domain (residues 352-433; *Syn*6803 number 343-424) that shares 45% protein sequence identity with *Syn*6803 RNAP SI3-tail domain (Fig. S4A and Table S1), and built a near-complete structure model of *Syn*6803 RNAP SI3, aided by our crystal structure of *Syn*7942 RNAP SI3-tail and a crystal structure of *Thermosynechococcus elongatus BP-1* RNAP SI3 (residues 435-983; *Syn*6803 residues 430-977; PDB: 8EMB).

The cyanobacterial RNAP-SI3 folds into a seahorse-shape structure that could be further divided into four domains: SI3-tail (residues 343-424), SI3-fin (residues 884-973 and 424-478), SI3-body (residues 478-619 and 729-884), and SI3-head (residues 619-729) (Fig. 2A). *Syn*6803 RNAP SI3-body and SI3-head domains are absent in *E. coli* RNAP SI3, while the SI3-tail and SI3-fin domains resemble the two domains of *E. coli* RNAP-SI3 (SI3-NTD and SI3-CTD) that are located near the secondary channel (Fig. 2B). The *Syn*6803 RNAP SI3-tail domain adopts a structural fold similar to the *E. coli* RNAP SI3-NTD [(root mean square deviation (RMSD) 1.7 Å of 296 Cα atoms; Fig. S4D)], while the SI3-fin domain of *Syn*6803 RNAP adopts a structural fold radically different from *E. coli* RNAP SI3-CTD (RMSD 3.6 Å of 364 Cα atoms). In contrast to the two sub-domains of *E. coli* RNAP-SI3 that stably associate with each other, the SI3-tail and SI3-fin domains of *Syn*6803 RNAP barely contact each other, which presumably permits independent movement of the SI3-tail (Figs. 2A and 2B). *Syn*6803 RNAP-SI3 contacts the rest of RNAP holoenzyme through multiple surface patches. SI3-tail and SI3-fin bind the loop and stem of the rim helices, respectively; SI3-fin contacts the β-lobe loop that stabilizes downstream dsDNA; SI3-body extends to the top of the main cleft and shields the lobe and protrusion domains; and the SI3-head makes interaction with σ^A^ (Figs. 2C and 2D).

**Figure 2.**
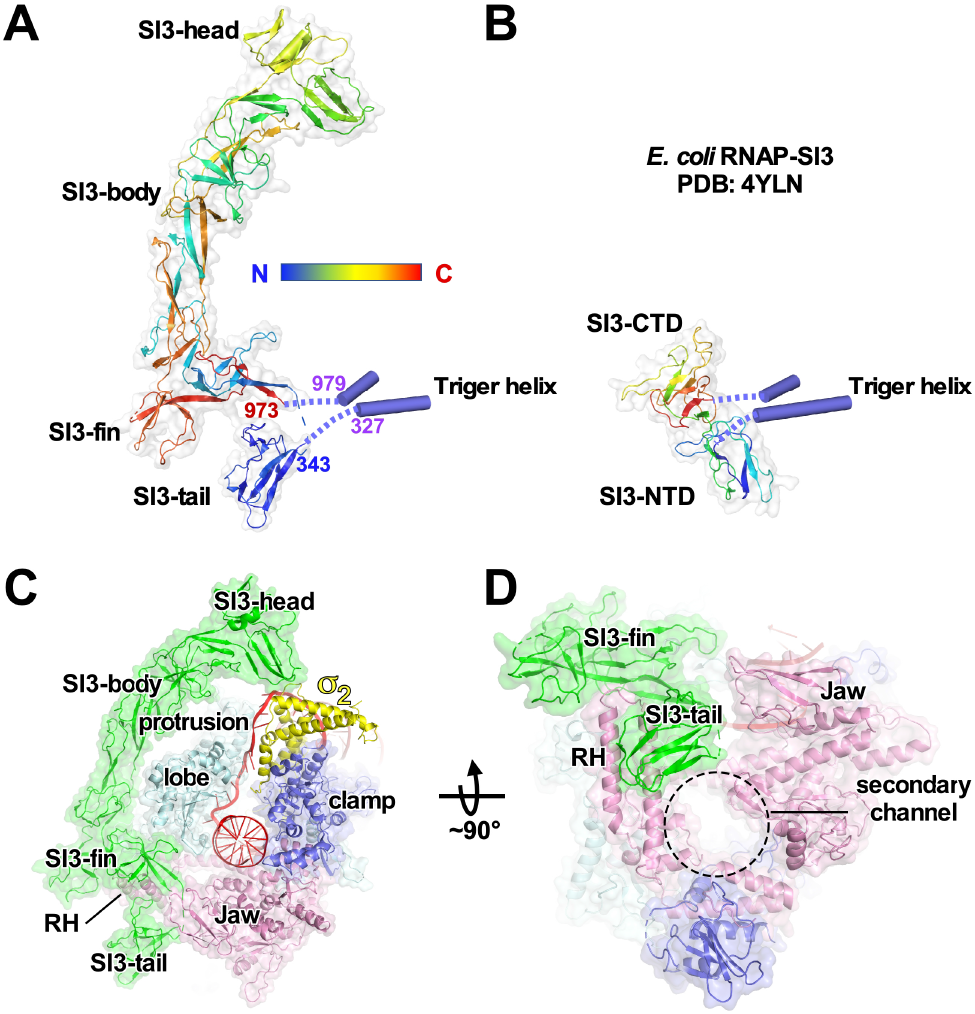
The structure and interaction of *Syn*6803 RNAP-SI3. **(A)** The overall structure of *Syn*6803 RNAP-SI3. The Trigger Helices connect to the SI3-tail and SI3-fin sub-domains. **(B)** The overall structure of *E. coli* RNAP-SI3 (PDB: 4YLN). **(C)** Overall structure shows that SI3 encloses half of RNAP surface. It extends from the secondary channel to the top of main cleft and makes interaction with σ_2_. **(D)** The SI3-fin and SI3-tail domains shield RNAP rim helixes (RH).

### Cyanobacterial RNAP-SI3 forms a SI3-σ arch with σ^A^

RNAP SI3-head tethers σ^A^ by making interactions with the σ^A^_1.2_ helix and the specificity loop of σ^A^_2_, a key structural motif that recognizes a flipped-out nucleotide within the −10 element of promoter DNA to initiate dsDNA melting. The SI3-σ interaction forms an arch-like structure that inhibits opening of the RNAP clamp and provides a physical obstacle that prevents the single-stranded DNA of the transcription bubble from dissociating and rewinding (Fig. 3A). Moreover, the negatively charged surface of the SI3-σ arch helps restrain the promoter DNA by electrostatic repulsion in the underneath positively charged groove (Fig. 3B). Although RNAP SI3 doesn’t contact promoter DNA, it fills in a shallow cavity between the specificity loop and σ^A^_1.2_ helix, likely stabilizing the active conformation of the specificity loop for accommodating the flipped −11A nucleotide of non-template strand promoter DNA (Figs. 3C and 3D).

**Figure 3.**
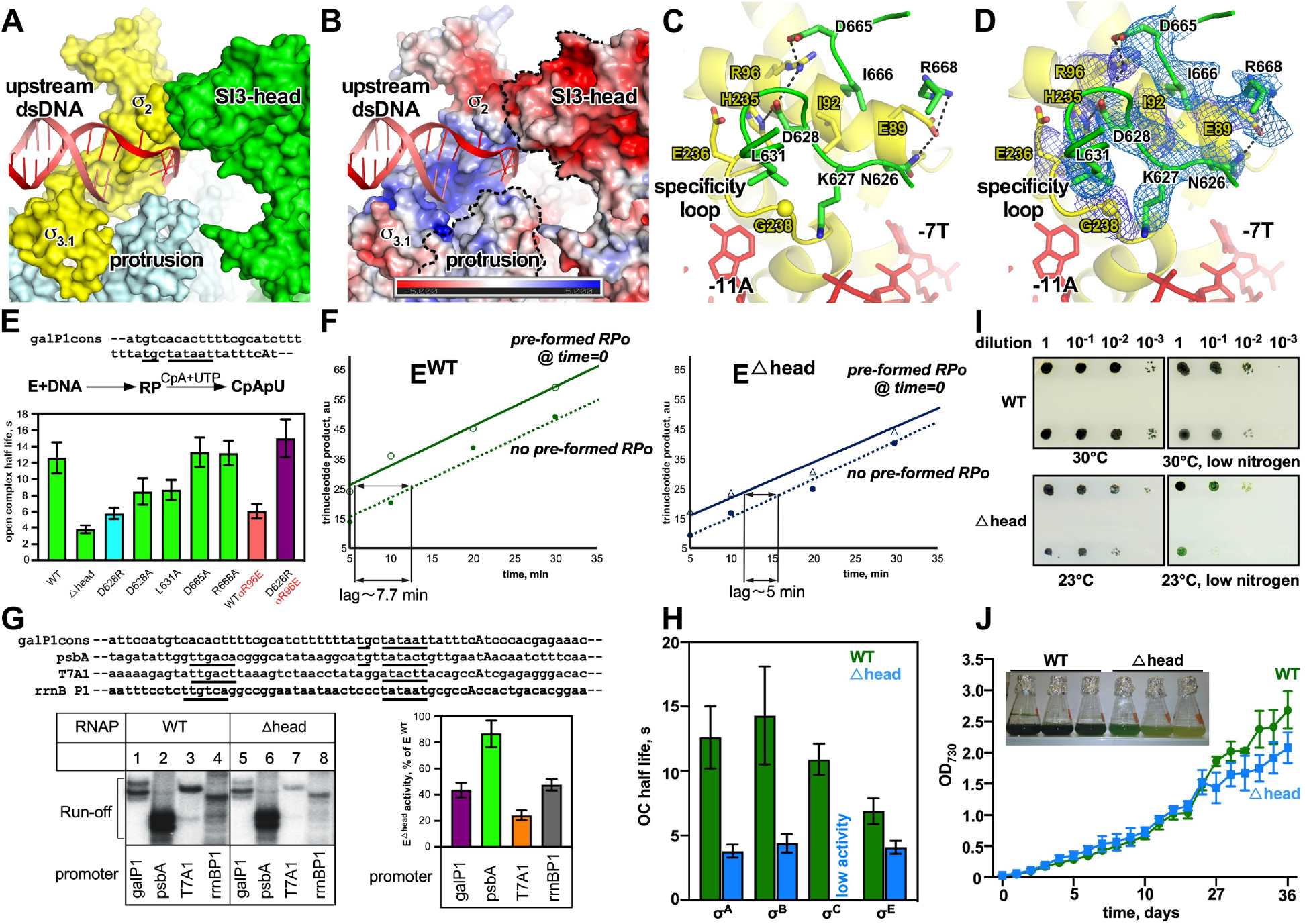
The structural and functional analysis of SI3-σ arch. **(A)** Surface presentation of the SI3-σ arch. **(B)** Electrostatic surface presentation of the SI3-σ arch. **(C)** Detailed interaction between SI3 and σ_2_. **(D)** The cryo-EM map of the SI3-σ_2_ interface. **(E)** The half-life of RNAP-promoter open complex comprising *gal*P1cons promoter and indicated holoenzymes. Error bars represent standard deviations from triplicate experiments. **(F)** Estimated time required for RPo complex formation of E^WT^ (left) and E^βhead^ (right). The timepoints are average of two independent experiments. **(G)** Activity of E^WT^ and E^βhead^ on different promoters. Top, promoter sequences with transcription start site shown in bold and consensus elements underlined; bottom, representative gel (left) and bar plot (right) shows amounts of run-off transcripts of E^βhead^ in % from that of E^WT^. Error bars represent standard deviations from 3 independent experiments. **(H)** The half-life of RNAP-promoter open complexes comprising WT/βhead RNAP and different σ factors. Error bars represent standard deviations from triplicate experiments. **(I)** Serial dilutions of cultures of *S. elongatus* 7942 WT strain and strain with genomic deletion of SI3 (βhead) plated on BG-11 media and grown at conditions indicated below images with constant light. **(J)** Growth curves of WT and Δhead *S. elongatus* 7942 strains in 12h light/12h dark conditions. Error bars represent standard deviation from three independently grown cultures. The insert is an image of flasks after 33-day growth.

Close interactions of the SI3-σ interface include two salt-bridge bonds made by D628 and D665 of SI3-head and R96 of σ^A^, one salt-bridge bond made by R668 of SI3 and E89 of σ^A^, one H-bond made by N626 of SI3-head and E89 of σ^A^, and one H-bond made by D628 of SI3-head and H235 of σ^A^. Moreover, residues K627, L631 and I666 of SI3 and residues I92, E236 and G238 of σ^A^ contribute to the interaction through Van der Waals force (Figs. 3C and 3D). The interface residues are conserved across different cyanobacteria species, suggesting physiological relevance of the SI3-σ interaction (Figs. S5A and S5B).

Contacts seen in cryo-EM structure between the SI3-head and σ^A^_2_ are expected to stabilize RNAP holoenzyme-promoter DNA complex. To test this hypothesis, we challenged transcription on a *gal*P1cons promoter with the DNA competitor heparin. We used WT *Syn*7942 RNAP holoenzyme along with mutants bearing a SI3 head domain deletion (E^Δhead^) or single amino acid substitutions of SI3 residues that make contacts with σ in the structure - E^D628A^, E^D628R^, E^L631A^, E^D665A^, and E^R668A^. The *gal* P1cons promoter, a model promoter of the extended −10 type that is predominant in cyanobacteria (31), was chosen to test the mutants. The RPo complexes were challenged with heparin (10 μg/mL) for increasing time intervals prior to addition of substrates for synthesis of the 3-nt RNA transcripts (Figs. 3E and S6A). The RPo half-life was calculated from the decay plots of the activity (Fig. S6A). Transcription reports directly on promoter complex half-life in this experiment, since steady-state rate of synthesis of the short abortive transcript is proportional to the amount of RPo complexes, and 10 μg/mL concentration of heparin does not affect catalysis of nucleotide addition. E^Δhead^ and E^D628R^ showed significant, up to 5 times decrease of the RPo half-life (Fig. 3E). Furthermore, the E^D628R^ effect was stronger than E^D628A^, highlighting the importance of the salt-bridge bond between D628 of SI3 and R96 of σ for stabilizing RPo (Figs. 3C and 3D). Notably, the impaired RPo stability of E^D628R^ was fully restored to WT level in E^D628R/σR96E^ holoenzyme, in which a reciprocal point amino acid change in σ, R96E was introduced to restore the salt-bridge bond (Fig. 3E). Control experiments showed that the mutant RNAP core enzyme (Δhead) exhibited the same catalytic rate and affinity to σ as those of the WT RNAP core enzyme (Fig. S7). Overall, these results validate interactions seen in the cryo-EM structure and highlight the importance of SI3-σ arch interaction on RPo stability.

The SI3-σ arch interaction might shift the dynamic conformational oscillation of the RNAP-β′ clamp/σ_2_ module. To test whether breaking of SI3-σ contacts increases the time required for promoter complex formation, we measured the kinetics of RNA production by E^WT^ or E^Δhead^ RNAP holoenzyme. Time interval needed for DNA to bind holoenzyme can be detected as a time lag between the moment after DNA addition and appearance of the RNA product. Time lag can be experimentally detected by comparing kinetics of the reaction started with simultaneous addition of DNA and NTPs, to a reaction where DNA was pre-incubated with holoenzyme prior to NTPs addition (32). In all reactions, RNA accumulates linearly over time, and there is a shift of the graph for reactions containing DNA-NTPs, reflecting time needed for DNA-RNAP complex formation (Fig. 3F). This time was indeed shorter for E^Δhead^, compared to E^WT^ (Fig. 3F). The slopes of the curves were similar, reflecting a similar rate of RNA synthesis by E^βhead^ and E^WT^. In short, the results suggest that SI3-σ arch slows RPo formation likely through sealing the main DNA left.

The above results show the SI3-σ arch increases RPo stability but slows down the RPo formation. To investigate the overall effect of SI3-σ arch on transcription, we measured the activity of the WT and E^βhead^ RNAP holoenzymes with representative promoters, including P_*psbA* (a strong promoter with near-consensus −35, extended −10, −10, and discriminator elements and an optimal 17-bp −35/-10 spacer), P_*gal* P1cons (extended −10 promoter), P_T7A1 (−35/-10 promoter), and P_*rrn*B P1 (−35/-10 promoter) (Fig. 3G). The results showed that disruption of SI3-σ arch interactions significantly decreased the activity of RNAP on P_T7A1, P_*rrn*B P1, and P_*gal* P1cons promoters but had little effect on the P_*psb*A (Fig. 3G). Since P_T7A1 and P_*rrn*B P1 are promoters that form short-lived RPo (33), these results suggest that the SI3-σ interaction is crucial for transcription activity of promoters that form unstable RPo. The input of SI3 deletion may depend additionally on other factors, such as a presence of a consensus −35 element, as in the case of P_*psb*A promoter.

Sequence alignment of σ^A^ with other alternative σ factors shows that the σ^A^ residues, which make interactions with SI3, are conserved in four other σ^70^-type group II σ factors in *Syn*6803 (σ^B^, σ^C^, σ^D^, and σ^E^), suggesting that SI3 likely makes interactions with these σ factors, and plays a role in transcription initiation by the alternative σ^70^-type factors (Fig. S5A). Supporting this prediction, the stability of RNAP-promoter DNA complex comprising E^βhead^ and σ^70^-type group II σ factors σ^B^, σ^C^, σ^E^ (RpoD2, 4 and 6 of Syn7942) was impaired to different extents compared with that of the respective WT RNAP holoenzymes (Fig. 3H). Stability of the open complex is lower for holoenzyme formed with σ^E^ even for E^WT^, consistent with changes of both Glu89 and Arg96 residues participating in formation of SI3-σ to Gln residues in this σ (Fig. S5A). Intriguingly, the SI3-σ interface does not appear to be conserved in plant chloroplast RNAP (Fig. S5B). As the SI3 domains of chloroplast RNAPs are larger in many plant species as compared with cyanobacteria, detailed structural information is required to see if SI3-σ arch preserved in chloroplast RNAP.

To investigate the physiological role of the SI3-σ arch, we constructed a chromosomal deletion of the SI3 head domain in *Syn*7942 and tested the growth phenotype of the resulting strain. Under optimal laboratory growth conditions, there was no obvious growth defect of this mutant on solid media at constant light (Fig. 3J). However, compromised growth of the SI3 head mutant strain was observed under stress-inducing conditions, such as low temperature, nitrogen deprivation, or their combination (Fig. 3I). In liquid culture with light conditions mimicking diurnal rhythms (12-hour light followed by 12-hour darkness), the WT and mutant strains initially grew at a similar rate, but the mutant strain entered stationary phase prematurely with bleached cultures after 20 days-growth (Fig. 3J). Altogether, these data suggest that SI3-σ arch plays a fundamental role in bacterial growth at nutrient limited conditions, as well as during a number of stress responses.

### Cyanobacterial RNAP-SI3 interacts with the rim helices

The cryo-EM structure of cyanobacterial RPitc reveals that SI3-tail makes extensive interactions with the rim helix hairpin near the secondary channel of RNAP (Figs. 2D, 4A, and 4B). The SI3-tail is a barrel-sandwich hybrid domain composed entirely of β strands. The ‘RTRHG’ loop (named after the five conserved resides of the loop) protrudes out from the main body of SI3-tail and contacts the stem of rim helix hairpin (Fig. 4A). The conserved residues R367, R369, H370, G371 of the ‘RTRHG’ make a H-bond network with four principal residues (R79, E89, K93, and N96) of the rim helix (Figs. 4A and 4B). The SI3-rim interface is conserved in all aligned cyanobacterial RNAP and chloroplast PEPs in various plants, but it is not conserved in *E. coli* and other bacterial species (Figs. 4D, S4C and S4D), suggesting a lineage-specific interaction. Disruption of this interaction by deletion of the RTHRG loop led to increased ubiquitous pausing during elongation and an inability for RNAP to reach the end of the template (Fig. 4C). This result suggests that disruption of SI3-rim interactions allows direct influence of SI3 movement on TL function at the active site (*see Discussion*).

**Figure 4.**
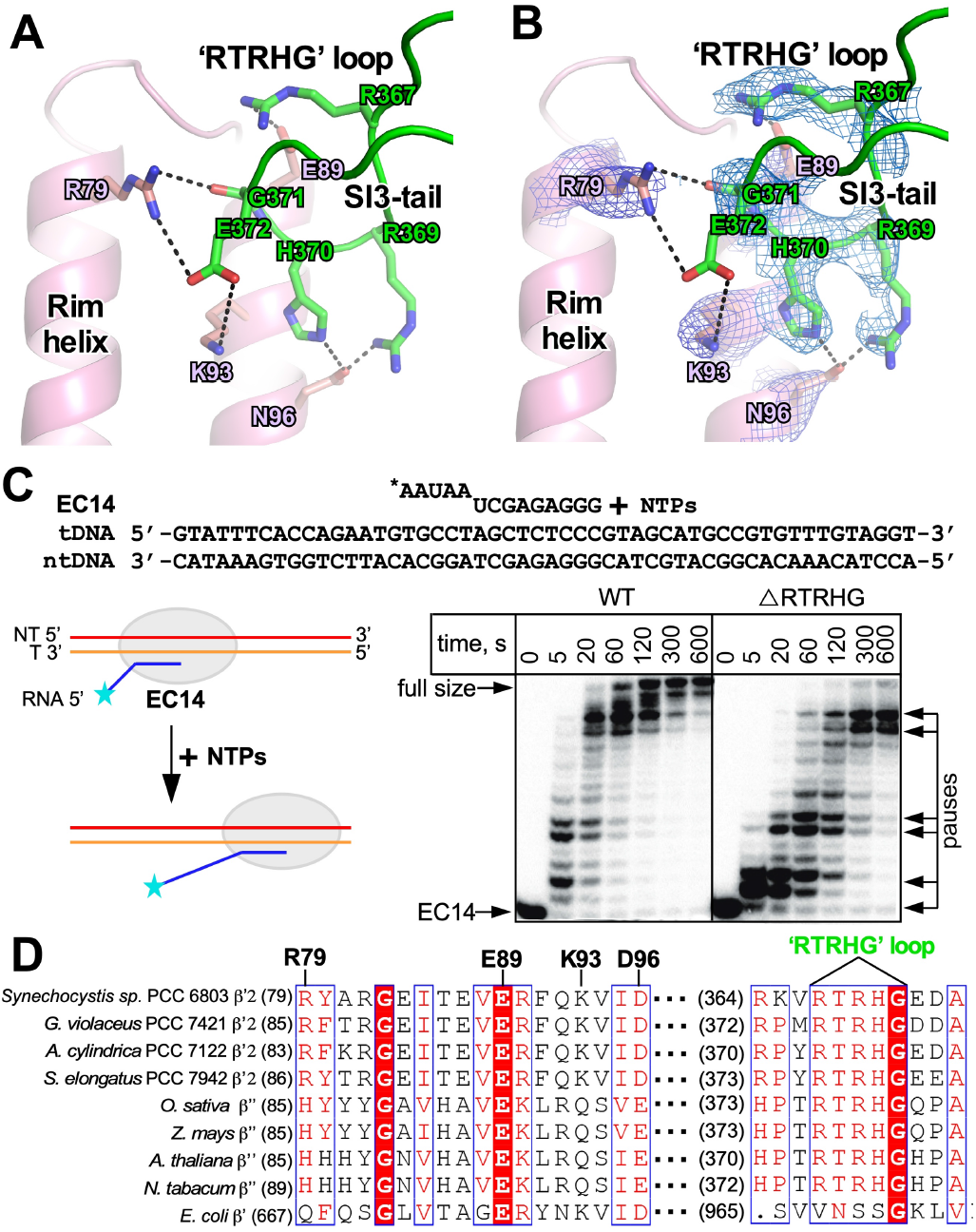
SI3 interacts with the rim helix. **(A)** Detailed interaction between the rim helices and SI3-tail. **(B)** The cryo-EM map of the SI3-Rim interface. **(C)** RTRHG loop deletion leads to increased pausing during elongation. Elongation complex is assembled with 14 nt long RNA labelled at 5’-end with ^32^P and template and non-template DNA oligonucleotides fully complementary to each other. NTPs were added to a final concentration of 10 μM, reaction stopped at indicated timepoints with addition of formamide-containing loading buffer. **(D)** The protein sequence alignment of SI3 tail of various cyanobacterial RNAP and plant chloroplast PEPs.

### The conformational change of SI3 upon TL refolding

The large SI3 insertion is located between the two helices of the TL (Fig. 2A), a key structural element that undergoes folding/unfolding during each nucleotide-addition cycle (16, 34). In the *Syn*6803 RPitc structure, the TL is in an unfolded state, probably due to absence of NTP at the ‘i+1’ site. To study whether refolding of TH affects SI3 conformation and its interaction with RNAP, we sought to determine the structure of *Syn*6803 NTP-bound RPitc. We first reconstituted *Syn*6803 RPitc with a modified RNA primer, where the 3’ terminal nucleotide was replaced by a 3’-deoxy adenine. We subsequently incubated the *Syn*6803 RPitc with CTP and determined the cryo-EM structure of *Syn*6803 CTP-bound RPitc at 3.0 Å resolution (Fig. S8). The cryo-EM map shows CTP occupies the ‘i+1’ site (Fig. 5A). α phosphate of CTP is in close distance with C3’ atom of the ‘i’ site adenine, indicating it adopts an ‘insertion’ state ready for incorporation (Fig. 5A).

**Figure 5.**
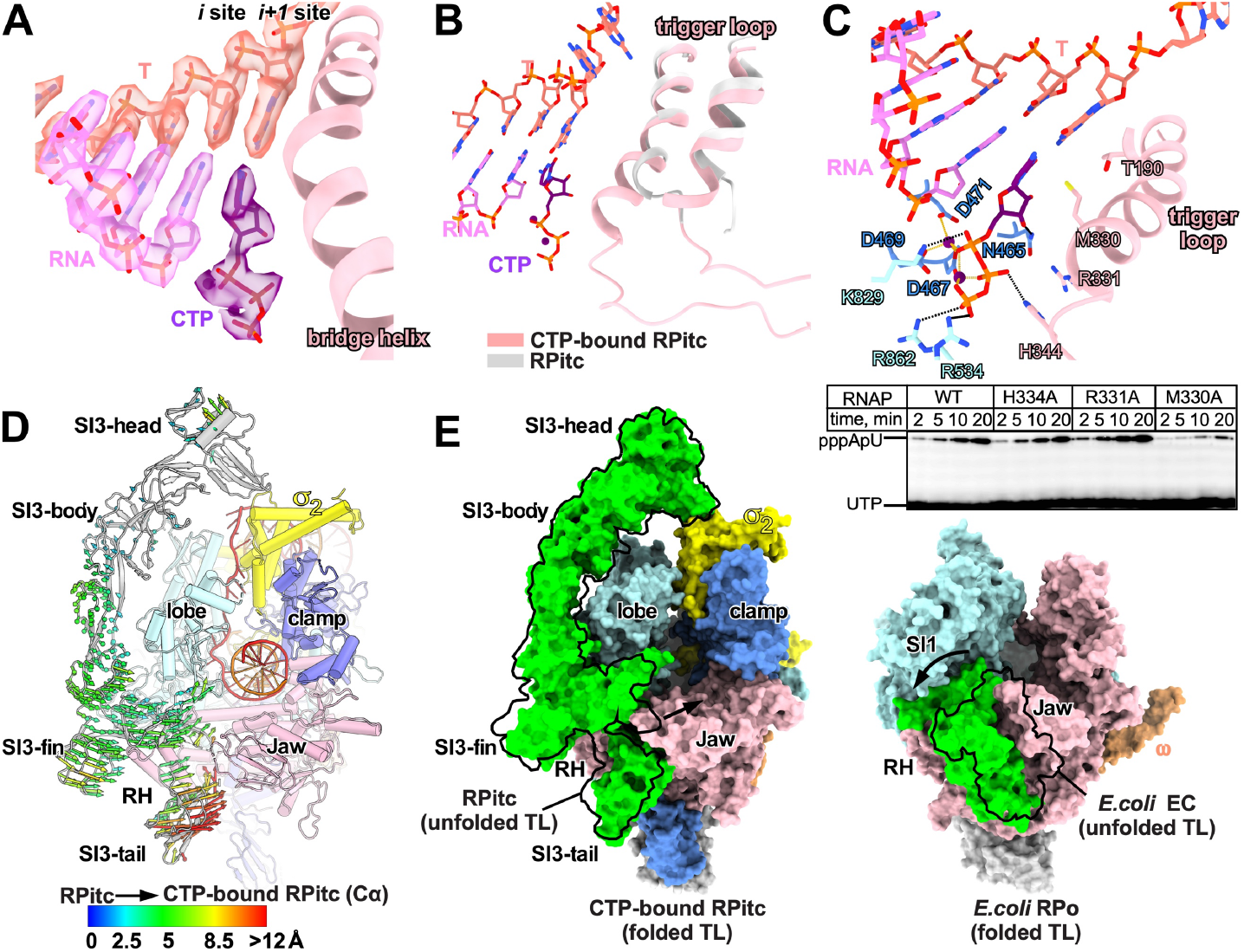
TL refolding induces structural change in Syn6803 NTP-bound RPitc. **(A)** The cryo-EM map and model for the active site. CTP adopts an ‘insertion’ state at the ‘i+1’ site. **(B)** CTP induces refolding of TL into TH. **(C)** The top panel shows the detailed interaction between CTP and RNAP residues; the bottom panel shows the effect of alanine substitutions of CTP-contact residues on the efficiency of the first phosphoric diester bond formation during *de novo* initiation on *gal*P1cons promoter. **(D)** The conformational changes of SI3 induced by TL refolding (cluster of arrows showing domain movement). **(E)** The comparison of SI3 movement upon TL refolding between *Syn*6803 RNAP (left) and *E. coli* RNAP (right).

CTP binding induces refolding of the TL into the TH (Fig. 5B). The refolded TH forms salt-bridge and H-bond interactions with the phosphate groups of CTP essentially the same as in the crystal structure of *T. thermophilus* CMPCPP-bound TEC (16), except that the invariant histidine (H334), which functions as a positional catalyst(13), contacts β phosphate in our structure instead of α phosphate in *T. thermophilus* CMPCPP-bound TEC (Fig. 5C and S9B). In agreement with the structure, mutant RNAP with an M330A change has a strong effect on the formation of the first diester bond between the initiating ATP and UTP on a *gal*P1cons promoter, presumably due to loss of its stacking interaction with the base of the incoming nucleotide (Fig. 5C). R331A and H334A substitutions have lesser effect (Fig. 5C).

Upon TL refolding, the conformation of the SI3-head and -body domains remain unchanged but SI3-fin and -tail domains undergo a rotational movement towards the secondary channel (Fig. 5D). The SI3-σ arch remains intact, suggesting nucleotide addition doesn’t disrupt the SI3-σ arch interaction (Fig. S9C). Interaction between ‘RTRGH’ of SI3-tail and β′ rim helices also remains intact, since β′ rim helices, SI3-fin, and SI3-tail rotate as a single structural unit (Figs. S9D and S9E). The TH refolding-induced stretching of the two short linkers, L1 and L2, that connect the TH to the SI3-tail and SI3-fin domains likely accounts for the domain rotation (Fig. S9F). Compared with the large conformational change of *E. coli* RNAP SI3 upon TH refolding (20, 35), this structural module in cyanobacteria RNAP rotates to a much lesser extent (Fig. 5E).

## DISCUSSION

Cyanobacteria are the only prokaryotes capable of oxygenic photosynthesis (36). As a result, they oxygenated the atmosphere of the Earth ∼ 2.3 Ga, changing the subsequent evolutionary course of the entire biosphere; they gave rise to chloroplasts ∼2.1 Ga (36, 37). Here we report the architecture of the cyanobacterial RNAP and pave the road for further understanding and engineering of the transcription apparatus in cyanobacteria. Moreover, since cyanobacterial RNAP is the ancestor of plastid-encoded RNAPs, our work provides foundation to understand the structure and evolution of chloroplast RNAP.

One of our intriguing findings is that the SI3 interacts with σ^A^ forming an arch-like structure on top of the main cleft that stabilizes the closed clamp of cyanobacterial RNAP. The SI3-σ arch could potentially affect at least two steps during transcription initiation. Firstly, the SI3-σ arch likely influences clamp dynamics of cyanobacterial RNAP holoenzyme, shifting the equilibrium towards the closed conformation. The closed SI3-σ arch presumably hinders loading of non-specific dsDNA into the main cleft and thereby could prevent pervasive transcription at non-promoter genomic regions. On the other hand, when the genuine promoters are recognized by cyanobacterial RNAP through sequence-specific interaction of −35/σ_4_ or extended −10/σ_3.1_, electrostatic repulsion between negatively charged DNA backbone and SI3-head domain could facilitate opening of the SI3-σ arch, which is glued only through a small interface (350 Å^2^) allowing loading of downstream promoter dsDNA (Fig. S10). Secondly, the SI3-σ arch can stabilize the RPo complex, as we showed here. After RNAP and promoter DNA form a catalytically competent RPo complex, the SI3-σ arch seals the top of main cleft to restrain the clamp in a closed conformation, and therefore prevents transcription bubble collapse.

We predict that the SI3-σ arch likely affects transcription initiation of a large proportion of genes, as the SI3-σ arch is present in majority of RNAP-σ holoenzymes, including alternative group 2 σ factors (Fig. S5A). However, the transcription outcome of a particular gene depends on the kinetics of transcription initiation formed at its promoter DNA. We have shown that an impaired SI3-σ arch decreases transcriptional output from representative promoters by different extents, which causes a pleiotropic effect on growth of cyanobacteria, especially in nitrogen depleted conditions. This pleiotropism could represent the cumulative effect of several σ factors involved, since in the absence of the specialized σ^54^ factor for nitrogen metabolism in cyanobacteria, this function is distributed between a number of σ factors, including σ^A^, σ^B^, σ^C^, σ^E^ (38-41). Further detailed study is required to globally evaluate the contribution of SI3-σ arch on cyanobacterial gene expression.

Folding of the TL upon substrate addition is completed without SI3-σ arch disruption or large conformational change of SI3 at the stage of transcription initiation, in contrast to *E. coli* SI3 (20). After RNAP escapes from the promoter and enters the elongation stage, the SI3-σ arch is broken, and SI3 “body” and “head” may become more mobile and potentially available for interactions with cellular factors. Could these interactions generate a signal for SI3 to transmit onto the TL, and affect catalysis? Perhaps not, at least not uncontrollably. We suggest that the TL is insulated from effects of SI3 conformational changes by SI3 attachment to the main body of RNAP at the base of the NTP entry channel. Disruption of this interaction via RTRHG loop deletion leads to increased pausing in elongation due to the effect of thermal motion of SI3 on TL folding. Moreover, this attachment and the lack of major conformation change in SI3 upon NTP binding implies a low probability of its rhythmic movement with every nucleotide addition cycle, like proposed movement of much smaller *E. coli* SI3 (42). At the same time, we can not exclude that, under specific conditions, the interface between Rim Helices and RTRHG loop could be targeted by regulatory factors to influence catalysis by the TL.

In summary, in this study we present the first structures of cyanobacterial transcription initiation complexes. The structures reveal an unexpected SI3-σ arch interaction that stabilizes RPo and maintains bacterial growth in nutrient-limited environments. Further structures of transcription elongation and termination complexes are required to illustrate the regulatory mechanism of cyanobacteria RNAP in these transcription stages.

## ACKNOWLEDGEMENT

This work was supported by the National Key Research and Development Program of China (2018YFA0900701) to Y.Z., Basic Research Zone Program of Shanghai (JCYJ-SHFY-2022-012) to Y.Z. Leverhulme Trust Research Grant RPG-2018-437 and EPSRC grant EP/N031962/1 to Y.Y. We thank Prof. Murakami Katsuhiko for kindly sharing the coordinate of *Thermosynechococcus elongatus BP-1* RNAP SI3, Dr. Liangliang Kong, Dr. Fangfang Wang, Dr. Guangyi Li, and Dr. Jialin Duan at the cryo-EM center of NFPS in Shanghai, Dr. Shenghai Chang at the cryo-EM center of Zhejiang University for cryo-EM data collection.

## AUTHOR CONTRIBUTIONS

Conceptualization: YZ, YY

Protein preparation: SL, GL, LY, MP, ZG

Data collection: LS, LY, JS, YF

Structure determination: LS, LY

Transcription assay: GL, MP

Funding acquisition: YZ, YY,

Supervision: YZ, YY, YF

Writing: YZ, YY

## Data and code availability

The cryo-EM map and coordinates were deposited in Protein Data Bank and Electron Microscopy Data Bank (Syn6803 RPitc: 8GZG and EMD-34397; Syn6803 CTP-bound RPitc: 8GZH and EMD-34398; Syn7942 RNAP SI3-tail: 8H02)

## STAR Methods

### Plasmids

The pET28a-TEV-*rpoD* was constructed by inserting the *Syn*6803 *rpoD* gene into pET28a-TEV plasmid using restriction sites NcoI and NotI. The pET28a-TEV-*Syn*7942 *rpoC2* SI3-tail was constructed by inserting the DNA fragment encoding *Syn*7942 RNAP SI3-tail (residues 350-433) into pET28a-TEV plasmid using restriction sites NcoI and NotI. The pACYCduet-*Syn*6803-*rpoAZ* was constructed by inserting the *Syn*6803 *rpoA, rpoZ* genes into the pACYCDuet (Merk Millipore) using restriction site pairs BamHI/NotI and NdeI/XhoI, respectively. The pETDuet-*Syn*6803-*rpoBC2* was constructed by inserting the *Syn*6803 *rpoB, rpoC2* genes into pETDuet plasmid (Merk Millipore) using restriction site pairs NcoI/NotI and NdeI/XhoI, respectively. The pETDuet-*Syn*6803-*rpoBC1C2*(3xGS) was constructed by inserting the *Syn*6803 *rpoC1* gene into pETDuet-*Syn*6803-*rpoBC2* through homologous recombination, resulting in a fused *rpoC1C2 peptide* by a 3xGS-linker.

*Syn7942* genes *rpoA, rpoB, rpoC1, rpoC2*, and *rpoZ* were PCR amplified from genomic DNA. Primers were design so as to add RBSs sequences before each gene and a 7-residues long poly-histidine tag at *rpoC2* C-terminus. After purification, the PCR products were ligated in pJET1.2/blunt (ThermoFischer Scientific) with T4 DNA Ligase (New England Biolabs) according to the manufacturer’s instructions. The resulting plasmids were named pJET-A, pJET-B, pJET-C1, pJET-C2, and pJET-Z respectively, and were used as template DNA for downstream PCRs. *RpoB* was PCRed and assembled onto pJET-A by Gibson Assembly to yield pJET-AB, then *rpoC1* was assembled onto pJET-AB to yield pJET-ABC1. *RpoZ* was assembled onto pJET-C2 to yield pJET-C2Z. *RpoC2* and *rpoZ* were then assembled onto pJET-ABC1 to yield pJET-ABC1C2Z. Finally, the operon containing all five genes was PCR amplified from pJET-ABC1C2Z and assembled onto plasmid pET28a to yield pET28a-SelRNAP.

To obtain pET28a-SelRNAP-ΔSI3head, plasmid pET28a-Syn7942 RNAP was PCR amplified with primers flanking the head of β’ SI3 domain. The resulting PCR product was re-circularized by Gibson Assembly without addition of any insert. All Gibson Assembly reactions were performed using NEBuilder® HiFi DNA Assembly Master Mix according to the manufacturer’s instructions. Briefly, 50 ng of backbone PCR were mixed with a two-fold molar excess of insert and 2.5 μL NEBuilder® HiFi DNA Assembly Master Mix in 5.0 μL final volume. The reaction was incubated at 50°C for 1 hour before transformation in NEB5α cells. Genes expressing His_6_-tagged RpoD1, RpoD2, RpoD4, and RpoD6 were cloned separately into pET28 expression vector. Detailed information of the plasmids and primers used in this study are listed in Tables S3 and S4.

### *Syn*6803 RNAP

The recombinant *Syn*6803 RNAP core enzyme was purified from *E. coli* BL21(DE3) (Novo protein, Inc.) cells carrying pACYCDuet-*Syn*6803-*rpoAZ* and pETDuet-*Syn*6803-*rpoBC1C2*(3xGS). Protein expression was induced at an OD_600_ of 0.6-0.8 by 1 mM IPTG at 18 °C for 16 h. Cells were lysed in lysis Buffer A (50 mM Tris-HCl, pH 7.7, 200 mM NaCl, 5% glycerol, 2 mM EDTA, 2 mM DTT, 0.1 mM phenylmethylsulfonyl fluoride (PMSF) and protease inhibitor cocktail (Biomake.cn. Inc.)) using an Avestin EmulsiFlex-C3 cell disrupter (Avestin, Inc.). The supernatant of lysate was precipitated by 0.6% (v/v) polyethylenimine (PEI). The pellet was collected and RNAP was extracted from the pellet with buffer 50 mM Tris-HCl, pH 7.7, 5% glycerol, 1 M NaCl, 2 mM DTT, and 2mM EDTA. The RNAP solution was further precipitated by addition of ammonium sulfate (final concentration; 29 g/mL). The pellet was collected and RNAP was extracted in NTA-binding buffer (20 mM Tris-HCl, pH 7.7, 5% glycerol, 400 mM NaCl, 5 mM β-mercaptoethanol). The supernatant was loaded on to a 5 mL column packed with Ni-NTA agarose (SMART, Inc.). The bound-RNAP was washed with 50 mL NTA-binding buffer containing 20 mM imidazole and eluted with Ni-NTA buffer containing 500 mM imidazole. The eluted fractions were mixed with TGED buffer (20 mM Tris-HCl, pH 7.7, 5% glycerol, 2 mM DTT, 2 mM EDTA) at ratio 1:1 and loaded onto a Mono Q column (MonoQ 10/100 GL, Cytiva) followed by a salt gradient of buffer A (20 mM Tris-HCl, pH 7.7, 200 mM NaCl, 5% (v/v) glycerol, 1 mM DTT) and buffer B (20 mM Tris-HCl, pH 7.7, 600 mM NaCl, 5% (v/v) glycerol, 1 mM DTT). The fractions containing target proteins were collected, concentrated to 5 mg/mL, and stored at −80 °C.

### *Syn*6803 σ^A^

The recombinant *Syn*6803 σ^A^ was purified in *E. coli* BL21(DE3) cells carrying pET28a-TEV-*rpoD*. The protein expression was induced with 0.4 mM IPTG at 18 °C for 16 h when OD_600_ reached 0.6-0.8. Cell pellet was lysed in lysis buffer B (50 mM Tris-HCl, pH 7.7, 500 mM NaCl, 5% (v/v) glycerol, 5 mM β-mercaptoethanol, and 0.1 mM phenylmethylsulfonyl fluoride (PMSF)) using an Avestin EmulsiFlex-C3 cell disrupter. The supernatant was loaded on to a 2 mL column packed with Ni-NTA agarose (Smart-lifesciences, Inc.). The bound proteins were washed by the lysis buffer B containing 20 mM imidazole and eluted with the lysis buffer B containing 400 mM imidazole. The eluted fractions were subjected to TEV protease cleavage while dialyzing to buffer 20 mM Tris-HCl, pH 7.7, 200 mM NaCl, 5% (v/v) glycerol, 5 mM β-mercaptoethanol. The sample was reloaded onto the Ni-NTA column to remove impurity. The sample was diluted, loaded onto a MonoQ column, and eluted with a salt gradient of buffer A (20 mM Tris-HCl, pH 7.7, 0.1 M NaCl, 5% (v/v) glycerol, 1 mM DTT) and buffer B (20 mM Tris-HCl, pH 7.7, 0.5 M NaCl, 5% (v/v) glycerol, 1 mM DTT). The fractions containing target proteins were collected, concentrated to 10 mg/mL, and stored at −80 °C.

### *Syn*7942 RNAP core enzyme and σ subunits

RNAP core enzyme and σ subunits were separately expressed in T7express cells (NEB). Briefly, the cultures were grown in LB at 37 °C until OD_600_ ∼0.6, transferred to 20 °C, induced with 1 mM IPTG, and further grown for 4 hours. Cells were collected, resuspended in lysis buffer (50 mM Tris HCl pH 8.0, 250 mM NaCl, 10% glycerol, 20 mM imidazole, 1 mM β-mercaptoethanol, and protease inhibitors cocktail (Roche, according to manufacturer’s instruction) and lysed by sonication, all steps were done at 4 °C. Soluble fraction was recovered by centrifugation for 15 minutes at 17000 g and applied to 5 ml HiTrap Q HP column (Cytiva). Fractions containing core subunits were eluted with lysis buffer containing 200 mM imidazole. Pooled eluate fractions were loaded onto pre-equilibrated Strep-Tactin XT gravity flow column. Column was washed with buffer (100 mM Tris HCl pH 8.0, 150 mM NaCl, 1 mM EDTA), protein eluted with same buffer containing 2.5 mM DTT. Fractions containing pure core enzyme (judged by SDS PAGE) were pooled, concentrated using Amicon Ultra 100 kDa cut-off centrifugal device, and dialysed overnight against storage buffer (40 mM Tris HCl pH 8.0, 200 mM KCl, 50% glycerol, 1 mM EDTA, 1 mM DTT). σ subunits were isolated using HiTrap Q HP and Superdex 200 chromatography (Cytiva), fractions were pooled, proteins were concentrated and dialysed against the same storage buffer. Mutant RNAP core enzyme and σ subunits were produced using side-directed mutagenesis and same purification steps as for WT proteins.

### *Syn*6803 RNAP-σ^A^ holoenzyme

*Syn*6803 RNAP core enzyme and *Syn*6803 σ^A^ were incubated in a ratio of 1:4 at 4 °C for 4 h. The mixture was applied to a Superdex 200 Increase 10/300 GL column (Cytiva) equilibrated in 20 mM Tris-HCl, pH 7.7, 150 mM NaCl, 1 mM DTT. Fractions containing *Syn*6803 RNAP holoenzyme was collected and concentrated to 10 mg/mL.

### *Syn*7942 RNAP SI3-tail

The *Syn*7942 RNAP SI3-tail were purified in *E. coli* BL21(DE3) cells carrying pET28a-TEV-*Syn*7942-SI3tail by a similar procedure as above except an additional purification step on a HiLoad 16/60 superdex 75 pg column (Cytiva). The protein in 10 mM Tris-HCl, pH 7.7, 50 mM NaCl, 1 mM DTT were collected and concentrated to 40 mg/mL.

### Nucleic-acid scaffolds

Nucleic-acid scaffolds for cryo-EM study of *Syn*6803 RPitc and CTP-bound RPitc were prepared from synthetic oligonucleotides (sequences in Figs. 1A and S8A) by an annealing procedure (95°C, 5 min followed by 2°C-step cooling to 25°C) in annealing buffer (20 mM Tris-HCl, pH 8.0, 200 mM NaCl).

### Crystal structure determination of *Syn*7942 RNAP SI3-tail

The initial screen of *Syn*7942 RNAP SI3-tail was performed by a sitting-drop vapor diffusion method. Crystals were grown from drops containing 1 μL 40 mg/mL protein and 1 μL reservoir solution (0.2 M ammonium acetate, 0.1 M Tris pH 8.5, 25% w/v polyethylene glycol 3,350) at 22 °C. Crystals were transferred into the reservoir solution containing 15% (v/v) (2R, 3R)-(−)-2,3-butanediol (Sigma-Aldrich) and flash-cooled in liquid nitrogen. Data were collected at Shanghai Synchrotron Radiation Facility (SSRF) beamline 19U1, processed using HKL2000 (43). The structure was solved by molecular replacement with Phaser MR using the structure of *E. coli* β’ SI3 (PDB: 2AUK) as a search model (44). Cycles of iterative model building and refinement were performed in Coot (45) and Phenix (46). The final model of *Syn*7942 RNAP SI3-tail was refined to Rwork and Rfree of 0.210 and 0.233.

### Cryo-EM structure determination of *Syn*6803 RPitc

*Syn*6803 RNAP holoenzyme and the nucleic-acid scaffolds were incubated in a ratio of 1:1.5 at room temperature for 15 min. The mixture was applied to a Superdex 200 Increase 10/300 GL column (Cytiva) equilibrated in 20 mM Tris-HCl, pH 7.7, 150 mM NaCl, 1 mM DTT. Fractions containing *Syn*6803 RP**itc** were collected and concentrated to 10 mg/mL. *Syn*6803 RPo was subsequently mixed with CHAPSO (Hampton Research, Inc.) to a final concentration of 8 mM prior to grid preparation. About 3.5 μL sample was applied onto the glow-discharged C-flat CF-1.2/1.3 400 mesh copper grids (Electron Microscopy Sciences) and the grid was handled and plunge-frozen in liquid ethane using a Vitrobot Mark IV (FEI) with 95% chamber humidity at 10 °C.

Data were collected on a 300 keV Titan Krios (FEI) equipped with a K2 Summit direct electron detector (Gatan). A total of 3432 images were recorded using the Serial EM in super-resolution mode with a pixel size of 0.507 Å, and a dose rate of 53.6 electrons/pixel/s. Movies were recorded at 250 ms/frame for 8 s (32 frames total) and defocus range was varied from −1.5 μm to −2.6 μm. Frames in individual movies were aligned using MotionCor2 (47), and contrast-transfer-function estimations were performed using CTFFIND4 (48). About 1,000 particles were picked and subjected to 2D classification in RELION 3.0 (49). The resulting distinct two-dimensional classes were served as templates and a total of 473,353 particles were picked out. The resulting particles were manually inspected and subjected to 2D classification in RELION 3.0 by specifying 100 classes. A 60 Å low-pass-filtered map was calculated from structure of *E. coli* RPo (PDB: 4YLN) as the starting reference model for 3D classification (35). A total number of 158,150 particles were used for constructing the final cryo-EM map. The final maps were further subjected 3D auto-refinement, CTF-refinement, Bayesian polishing, and post-processing in RELION 3.0 (Fig. S3). Gold-standard Fourier-shell-correlation analysis (FSC) indicated a mean map resolution of 3.14 Å. The x-ray structure of *E. coli* RPo (PDB: 4YLN) was manually fit into the cryo-EM density map using Chimera. Model building and real-space refinement were performed in Coot (45) and Phenix(46).

### Cryo-EM structure determination of *Syn*6803 CTP-bound RPitc

*Syn*6803 RNAP core enzyme and σ^A^ were incubated in a ratio of 1:4 at 4 °C temperature for 4 h. The mixture was applied to a Superdex 200 Increase 10/300 GL column (Cytiva) equilibrated in 10 mM Hepes, pH 7.5, 100 mM KCl, 5mM MgCl_2_, 3 mM DTT. Fractions containing *Syn*6803 RNAP holoenzyme were collected and concentrated to 14 mg/mL. *Syn*6803 RNAP holoenzyme (30 μM) was mixed with the nucleic-acid scaffold (45 ωM) with a molar ratio of 1:1.5 for 1 h, and subsequently supplemented with CTP (3 mM) allowing incubation at 4 °C temperature for 4 h. The CTP-bound RPitc was mixed with CHAPSO (Hampton Research, Inc.) to a final concentration 8 mM prior to grid preparation. About 3 μL mixture was applied on a glow-discharged UltraAuFoil R1.2/1.3 300 mesh holey Au grids (Quantifoil Micro Tools GmbH), blotted with Vitrobot Mark IV (FEI), plunge-frozen in liquid ethane with 100% chamber humidity at 22 °C.

The micrographs were collected using EPU in the super-resolution counting mode on a 300 keV Titan Krios (FEI) equipped with a Gatan K3 Summit direct electron detector (pixel size 1.10 Å/pixel). A total of 2,297 images were recorded using the Super-resolution mode (exposure, 2.69 s per 40-frame movie; dose rate, 22.5 electrons/pixel/s; defocus, −1.2 to −2.2 ωm). Frames in individual movies were aligned using MotionCor2 (47), and contrast-transfer-function estimations were performed using CTFFIND4 (48). About 1,000 particles were picked and subjected to 2D classification in RELION 3.0 (49). The resulting distinct two-dimensional classes were served as templates and a total of 681,334 particles were picked out.

The particles were subjected to 2D classification in RELION 3.0 by specifying 100 classes. A 50 Å low-pass-filtered map was calculated from cryo-EM map of *Syn*6803 RPitc as the starting reference model for 3D classification (N=6). One 3D class with distinct shape of RNAP containing 120,655 particles was subjected to 2D classification again for generating templates for the second round of particle auto-picking. A total of 338,818 particles were auto-picked by using the 2D references. The particles were subjected to 2D classification in RELION 3.0 by specifying 100 classes. A total of 264,653 particles were selected and subjected to 3D classification using a 50 Å low-pass-filtered cryo-EM structure of the previous 3D classification (N=4). One 3D class with distinct shape of RNAP containing 145,731 particles were used for constructing the final cryo-EM map. The final maps were further subjected 3D auto-refinement, CTF-refinement, Bayesian polishing, and post-processing in RELION 3.0. Gold-standard Fourier-shell-correlation analysis (FSC) indicated a mean map resolution of 2.96 Å. The structure of *Syn*6803 RPitc was manually fit into the cryo-EM map using Chimera. Model building and real-space refinement were performed in Coot (45) and Phenix(46).

### Growth and mutagenesis of *Syn*7942

Wild type and mutant strains of *Syn7942* were grown in BG-11medium, at either constant light or 12-hour light/12-hour dark cycle as indicated in the legend of Figure 3 at 100 µE light intensity at 30 °C unless overwise indicated, in AlgaeTron 130 incubator. Liquid cultures were shaken at 300 rpm. Solid media was prepared using 1.2% gellan gum. Nitrogen starvation was induced by lowering down the concentration of NaNO_3_ in media to 10 µM.

To generate the mutant *S. elongatus* strains β’ΔSI3head and β’WT, the corresponding editing plasmids pUC-β’ΔSI3head and pUC-β’WT were transformed in wild type *S. elongatus* cells as previously described (50). Briefly, 15 mL of *S. elongatus* culture at OD_730_ 0.6-0.7 were pelleted by centrifugation at 6,000 g for 10 minutes. Cells were then washed once in 10 mL 10 mM NaCl and resuspended in 0.3 mL BG-11 medium. After addition of 0.5-5.0 µg of plasmid, cells were incubated overnight at 30 °C in the dark with gentle shaking and finally seeded on nitrocellulose membranes (purchased from Merck) laid on BG-11 agar plates supplemented with selective antibiotic. The plates were incubated under constant light at 30 °C and the membranes were transferred to fresh BG-11 agar plates every 2-4 days until colonies appeared. Colonies were then re-streaked several times until full segregation of the mutation was achieved (as assessed by PCR).

### *In vitro* transcription

All reactions for *Syn7942* RNAPs were done at 30 °C on linear PCR-derived templates containing promoters indicated on figures, in transcription buffer (20 mM Tris-HCl pH 7.9, 40 mM NaCl, 10 mM MgCl_2_). Reactions contained 30 nM RNAP core enzyme, 100 nM σ, and 100 nM promoter fragment. For testing susceptibility of promoter complexes to heparin treatment, 0.01 mg/mL heparin was added for 20”, 1’, 2’, 5’, 10’ before substrates addition. Transcription was initiated with 100 μM CpA and 40 μM [α-^32^P] radiolabeled UTP (7.5 Ci/mmol). Reactions were allowed to proceed for 5 min and were terminated by the addition of an equal volume of loading buffer (1X TBE, 8M Urea, 20 mM EDTA, 100 μg/mL heparin, 0.02 % bromophenol blue, 0.02 % xylene cyanole in formamide). Reaction products were resolved by electrophoresis in 23% denaturing polyacrylamide gel, visualized by PhosphorImager (Cytiva), and quantified using the ImageQuant software (Cytiva). Half-life of the promoter complex was calculated by fitting the data into exponential decay equation f = y_0_+a*exp(−b*x) using non-linear regression by SigmaPlot software, where x is the reaction time, f is the quantified 3-nt transcripts, y_0_, a, and b are unconstrained constants.

Reaction of dinucleotide product formation on *gal*P1cons promoter were performed with 500 µM ATP and 25 µM [α-^32^P] UTP (5 Ci/mmol), kept for the time intervals indicated in Figure, stopped with formamide-containing loading buffer and resolved in 33 % denaturing polyacrylamide gel. For testing the activity of RNAP on different promoters, the run-off transcription reactions contained 250 µM ATP, CTP, GTP and 25 µM [α-^32^P] UTP (5 Ci/mmol) were incubated for 10 minutes before termination with loading buffer.

For estimating the time intervals for RPo formation, transcription reactions were performed in 10 μL transcription buffer. Reactions contained 100 nM core RNAP (WT or Δhead), 300 nM σ^A^ and 50 nM promoter DNA. For control reactions components were incubated at 30° C 10 minutes to allow formation of open complexes. Reactions started by addition of 50 µM CpA di-nucleotide, 2 µM UTP [α-^32^P]-UTP (25 Ci/mmol) and were allowed to proceed for various amounts of time (for most reactions for 5”, 10”, 20”, 30” and 60”) at 30 °C before stopping by adding equal volume of stop buffer. For the test reactions 100 nM DNA was mixed with the same amount of CpA and [α-^32^P]-UTP in transcription buffer. The holoenzyme complex (WT/ Δhead core RNAP and σ^A^, pre-incubated for 10’ at 30 °C) was added to the DNA and substrates mixture to start reactions for the same amounts of time and stopped using equal volumes of formamide-containing loading buffer. RNA products were resolved in denaturing 23% acrylamide gels and visualised and quantified as before. Kinetics of RNA accumulation were fitted into linear equation y=y0+ax using linear regression by SigmaPlot software, where x is the reaction time, y is the quantified 3-nt transcripts, x is reaction time, y_0_ and a are unconstrained constants. Time interval for open complex formation was calculated by subtracting x value of test from control curve at y=0.

Transcription elongation kinetics experiments were performed using assembled elongation complexes essentially as described in (26). Briefly, elongation complex is assembled with 14 nt long synthetic oligonucleotide RNA labelled at 5’-end with [γ-^32^P] ATP and fully complementary template and non-template DNA oligonucleotides, sequences of all shown on Figure 4C. Assembled complexes were immobilized on Ni-NTA Sepharose beads via hexa-histidine tag on RNAP and washed with 1M NaCl-containing transcription buffer to remove any aberrantly assembled complexes. NTPs were added to final concentration of 10 ωM, reaction started with addition of 10 mM MgCl_2_ and stopped at timepoints indicated on Figure 4C with addition of formamide-containing loading buffer. Reaction products were resolved and visualized as before.

### Microscale thermophoresis

σ^A^ was fluorescently labelled on amines with NT647 RED-N-hydroxysuccinimide (NHS) reactive dye (Nanotemper), according to manufacturer’s protocol (Nanotemper). A fixed concentration of 400 nM σ^A^ was used in a set of 16 serial dilutions testing a range of core RNAP concentrations (from 0.5 nM to 5 µM) in MST buffer (40 mM Tris-HCl pH 7.9, 20 mM KCl, 10 mM MgCl2, 5% glycerol). 5 μL of each reaction was loaded into premium capillaries, and MST was performed at 80% fluorescence excitation power on a Monolith NT.115. Binding curves were plotted and *K*_*d*_ estimated using NT Analysis 1.5.41 and Affinity Analysis software.

## SUPPLEMENTAL TABLES

**Table S1.**
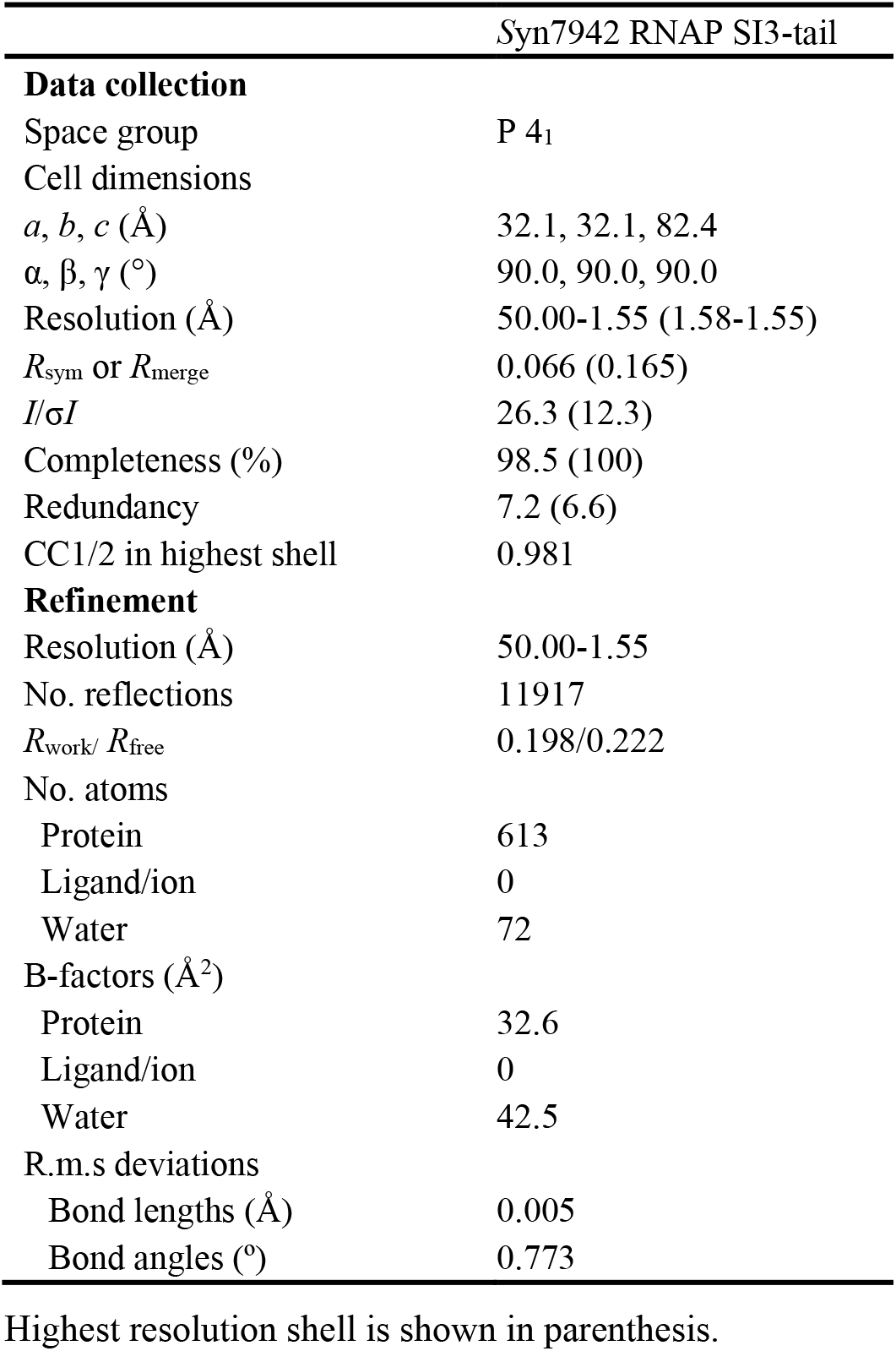
The statistics of *Syn*7942 RNAP SI3-Tail crystal structure.

**Table S2.** The statistics of *Syn*6803 RPitc cryo-EM structure.

|  | *Syn6803*<br>RPitc | *Syn6803*<br>CTP-bound RPitc |
| --- | --- | --- |
| **Data collection** |  |  |
| Number of grids used | 1 | 1 |
| Grid type | C-flat | UltrAuFoil |
| Microscope/detector | Titan Krios/Gatan K2 | Titan Krios/Gatan K3 |
| Voltage (keV) | 300 | 300 |
| Dose rate (e<sup>-</sup>/s) | 6.7 | 22.5 |
| Pixel size (Å/pix) | 1.014 | 1.1 |
| Total dose (e<sup>-</sup>/ Å<sup>2</sup>) | 53.6 | 50 |
| Total exposure time (s) | 8 | 2.69 |
| Number of frames/movie | 32 | 40 |
| Defocus range (µm) | -1.5 to -2.6 | -1.2 to -2.2 |
| Number of micrographs | 3,432 | 2,297 |
| Particles used for final map | 158,150 | 145,731 |
| **Model composition** |  |  |
| Non-hydrogen atoms | 29,547 | 30,339 |
| Protein residues | 3,630 | 3,693 |
| Nucleotide | 99 | 104 |
| Ligands (Zn<sup>2+</sup>/Mg<sup>2+</sup>/CTP) | 2/1/NA | 2/2/1 |
| **Refinement** |  |  |
| Resolution (Å) | 3.14 | 2.96 |
| Map sharpening B factors | 64.01 | -65.84 |
| Clash score | 3.31 | 3.46 |
| Average B factor (Å<sup>2</sup>) |  |  |
| Protein | 20.39 | 16.61 |
| Nucleotide | 39.50 | 31.58 |
| Ligand | 26.99 | 19.25 |
| RMS deviations |  |  |
| Bond lengths (Å) | 0.003 | 0.004 |
| Bond angles (°) | 0.592 | 0.762 |
| Ramachandran plot |  |  |
| Favored (%) | 96.28 | 95.44 |
| Allowed (%) | 3.72 | 4.56 |
| Outliers (%) | 0.00 | 0.00 |

**Table S3.** Constructs used in this study.

| Constructs | Sources |
| --- | --- |
| pET28a-TEV-*Syn6803-rpoD* | This study |
| pET28a-TEV-*Syn7942*-RNAP-SI3-tail | This study |
| pACYC-*Syn6803-rpoAZ* | This study |
| pCOLA-*Syn6803-rpoBC1C2(3xGS)* | This study |
| pJET-A | This study |
| pJET-B | This study |
| pJET-C1 | This study |
| pJET-C2 | This study |
| pJET-Z | This study |
| pJET-AB | This study |
| pJET-ABC1 | This study |
| pJET-C2Z | This study |
| pET28-*Sel7942*-rpoABC1C2Z | This study |
| pET28-*Sel7942*-rpoD1 | This study |
| pET28-*Sel7942*-rpoD2 | This study |
| pET28-*Sel7942*-rpoD4 | This study |
| pET28-*Sel7942*-rpoD5 | This study |
| pET28-*Sel7942*-rpoD6 | This study |
| pUC19-*Sel7942*-rpoC2/Spec/HR | This study |
| pUC19-*Sel7942*-rpoC2Δhead/Spec/HR | This study |

**Table S4.**
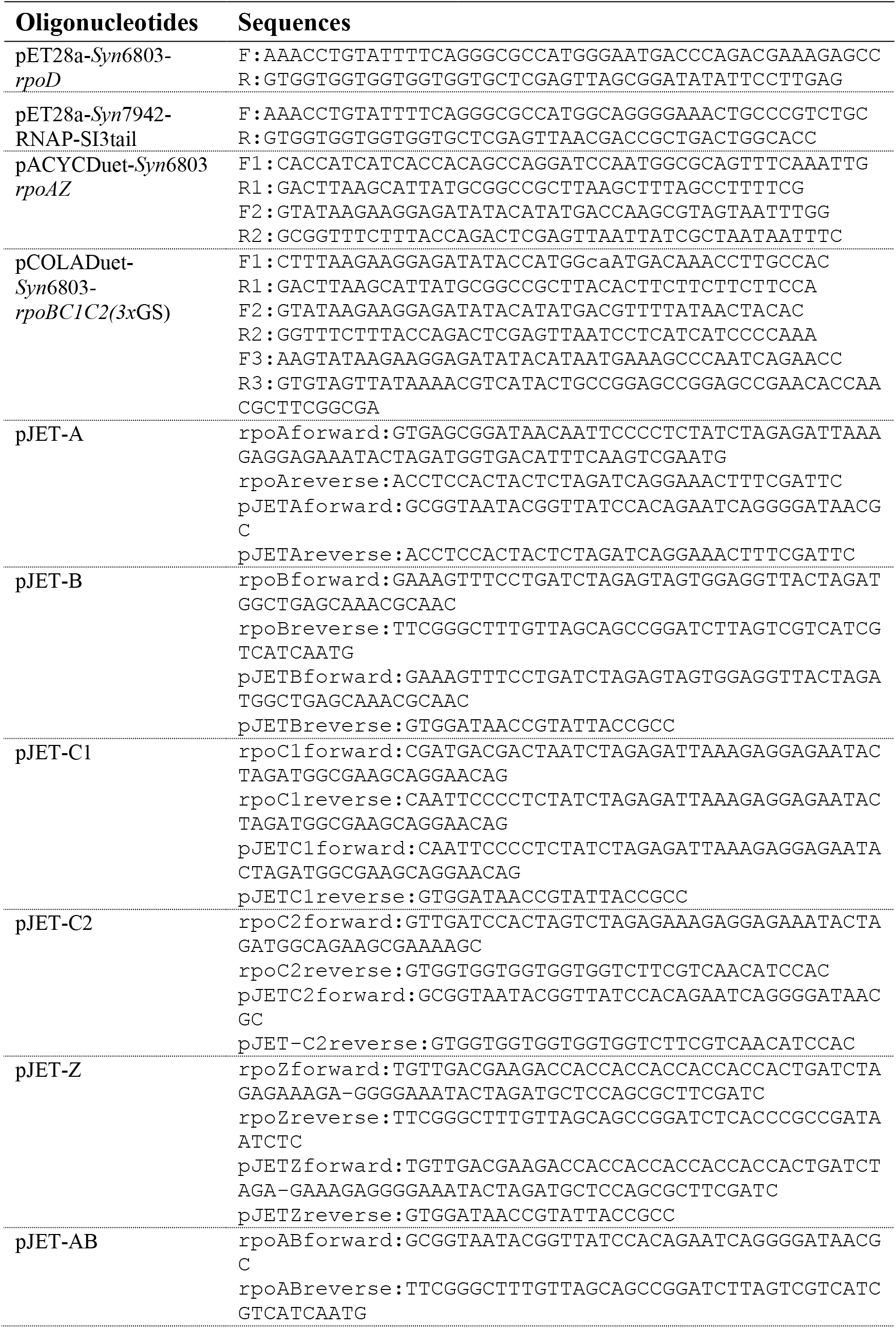
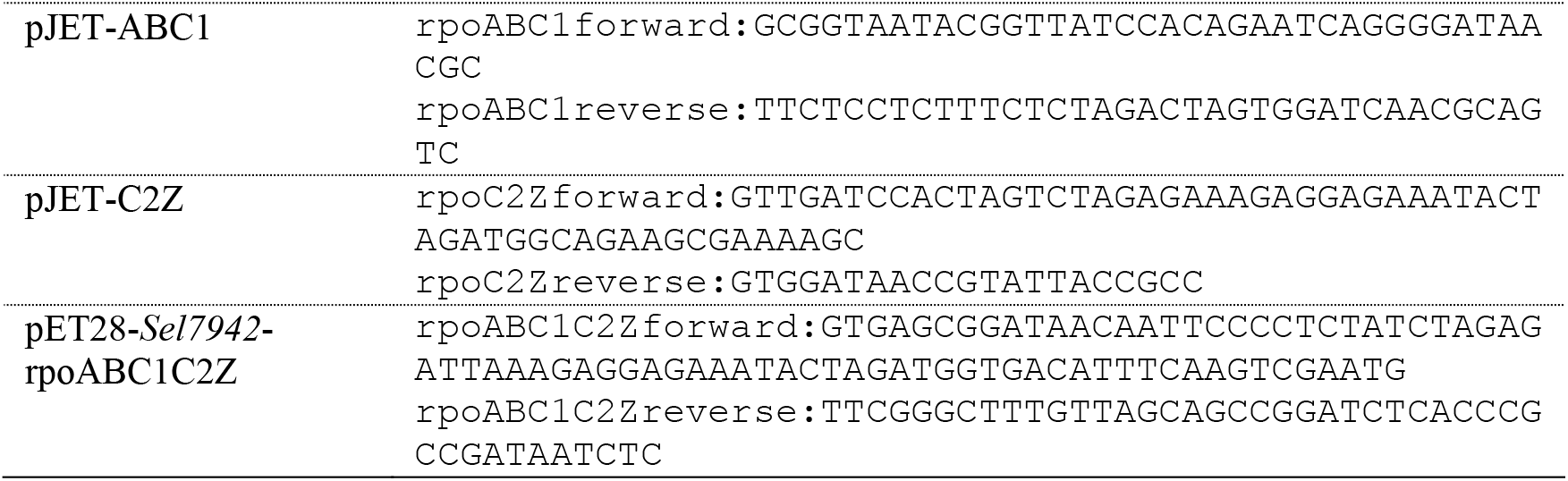
The primers sequences (5’ to 3’) used in this study.

## SUPPLEMENTAL FIGURES

**Figure S1.**
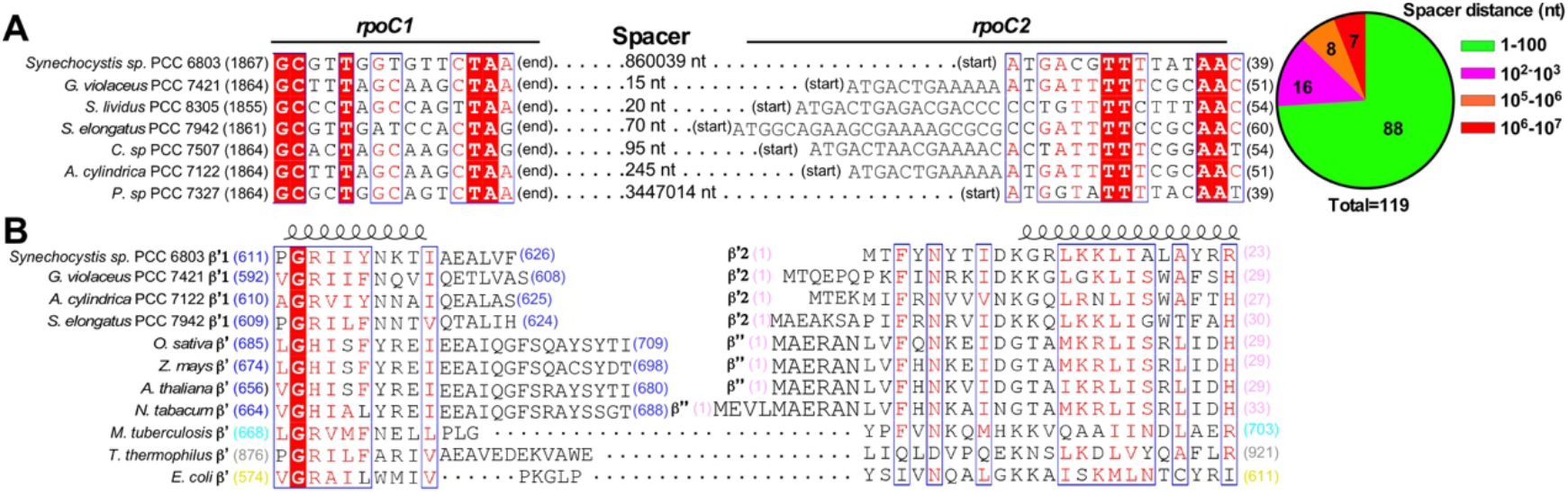
Sequence alignment of RNAP subunits of various cyanobacteria. **(A)** The DNA sequence alignment of *rpoC1* and *rpoC2* genes from representative cyanobacteria. The pie chart at right shows the spacer distribution of the two genes at 119 non-redundant cyanobacteria genomes. **(B)** The protein sequence alignment of RNAP-β’1 and -β’2 subunits from representative cyanobacteria, plastid-encoded polymerase (PEP) β’ and β’’ subunits of representative plant chloroplasts, and other eubacteria species.

**Figure S2.**
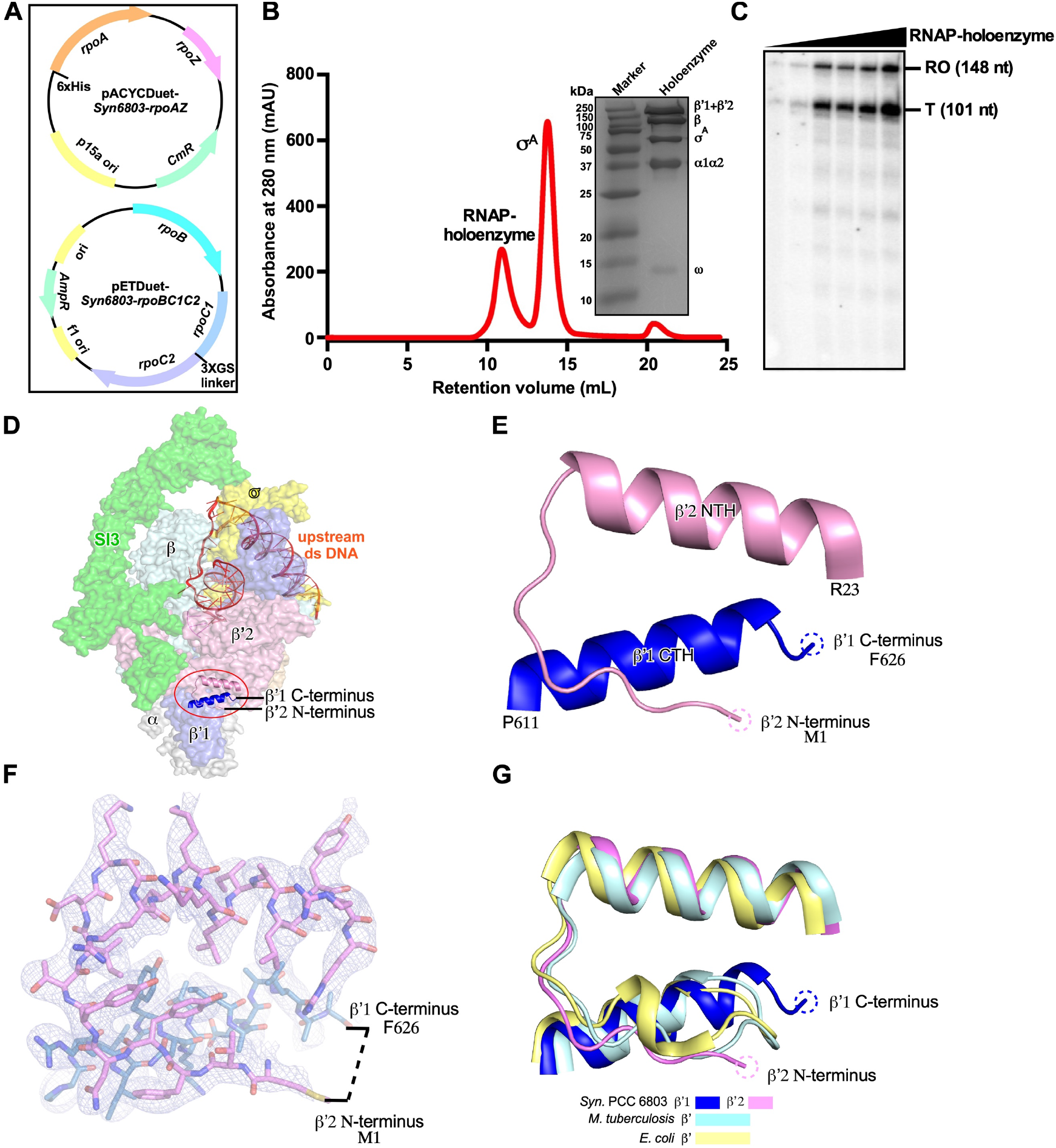
The assembly of *Syn*6803 RNAP-σA holoenzyme. **(A)** The schematic diagram of constructs used for over-expression of *Syn*6803 RNAP core enzyme in *E. coli*. **(B)** The gel-filtration chromatography result of RNAP-σA holoenzyme assembly, the SDS-PAGE shows the purity of *Syn*6803 Rpitc. **(C)** The *in vitro* transcription result of *Syn*6803 RNAP-σA holoenzyme using N25 promoter. RO, run-off product; T, terminated product. **(D)** The split occurs at the surface of RNAP. **(E)** The split ends remain close to each other. **(F)** The cryo-EM map (mesh) of the split ends. **(G)** structural superimposition of bacterial RNAP suggests that the split ends retain conserved structural fold.

**Figure S3.**
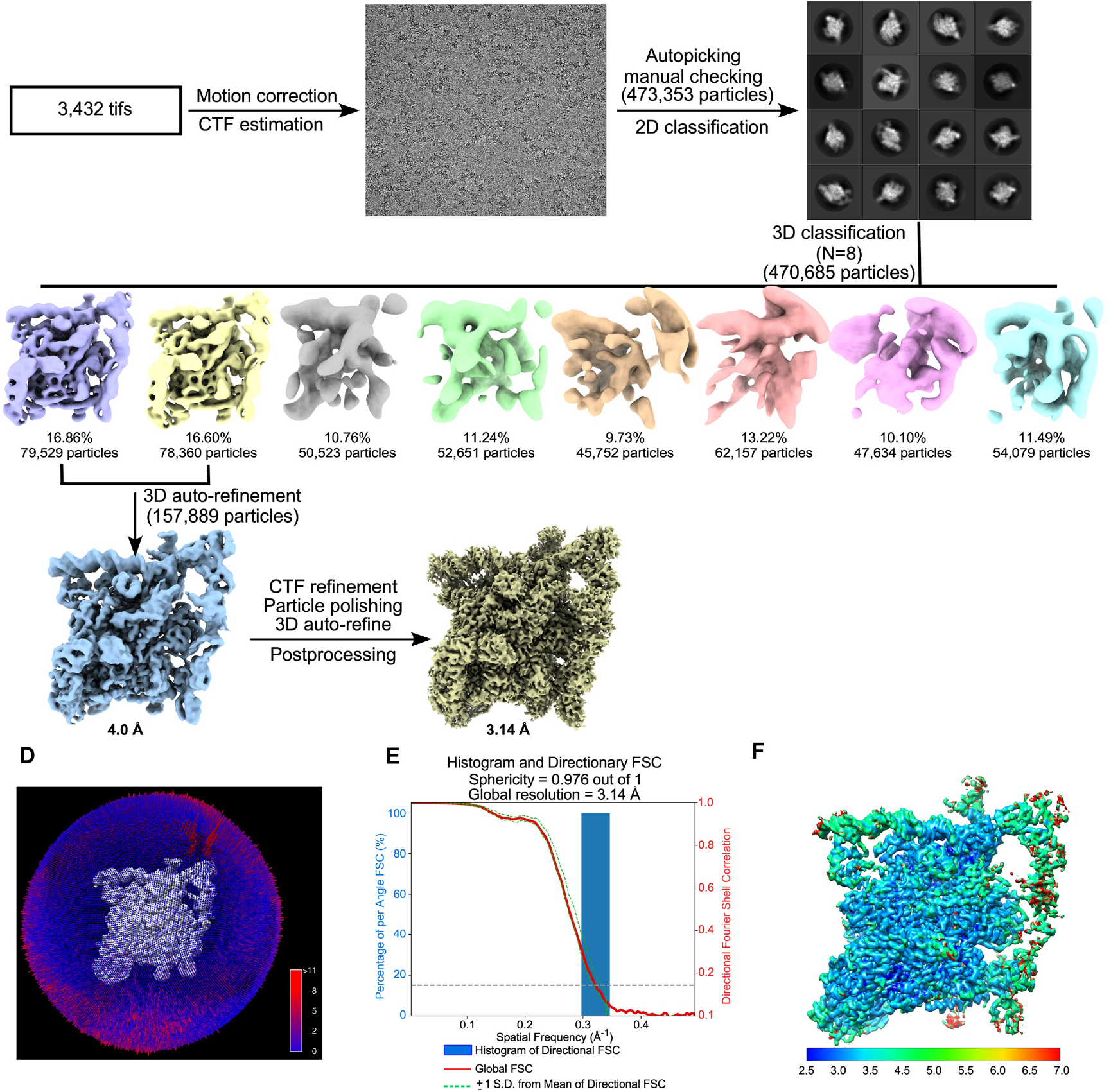
The flow chart of cryo-EM data collection and map calculation for RPitc. **(A)** The flowchart of data processing. **(B)** The angular distribution of single-particle projections by number of particles of each projection. **(C)** The 3D FSC plot. The dotted line represents 0.143 cutoff of the global FSC curve. **(D)** The cryo-EM map of *Syn*6803 RPitc colored by local resolution.

**Figure S4.**
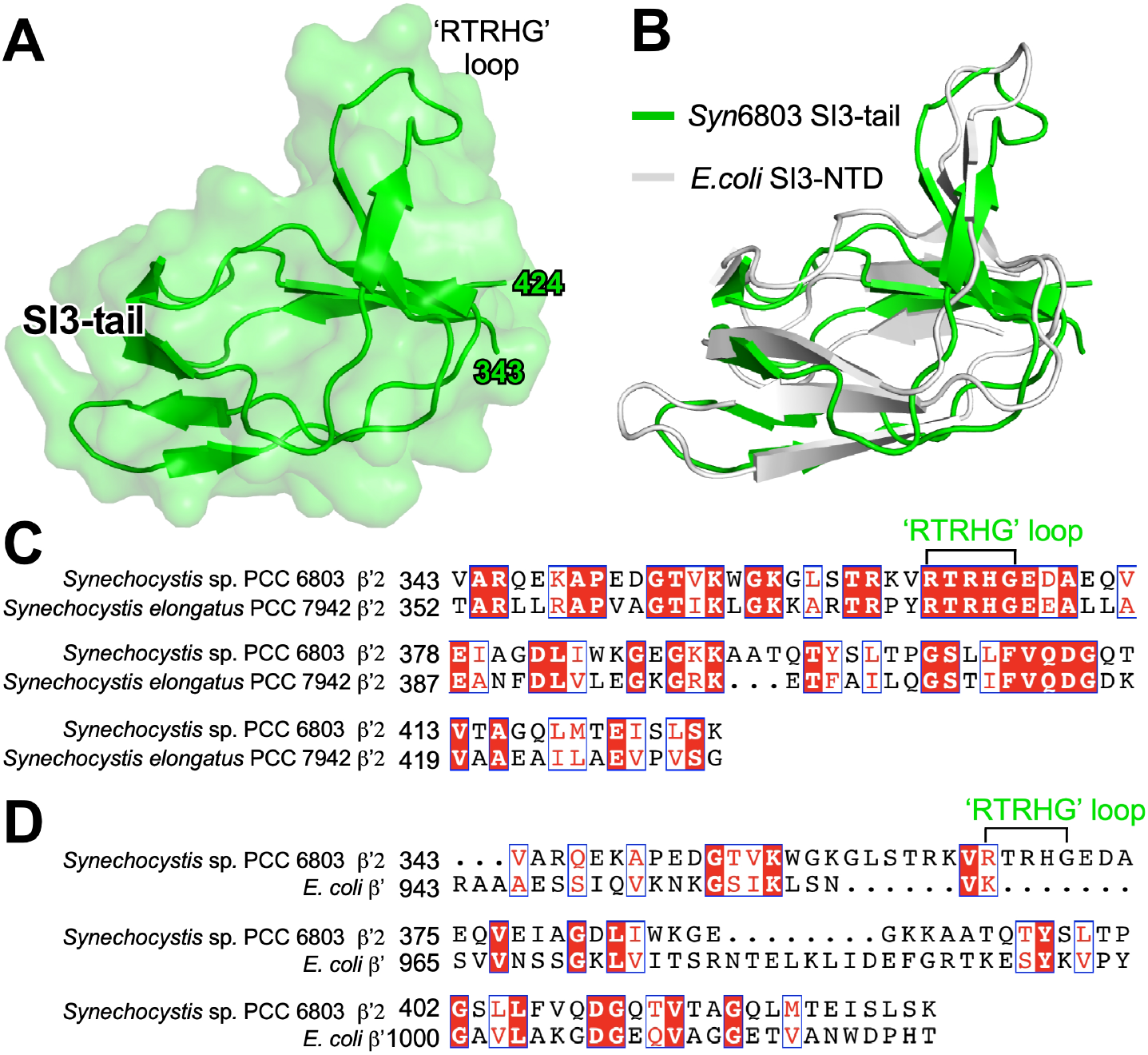
The crystal structure of *Syn*7942 RNAP SI3-tail. **(A)** The overall crystal structure of *Syn*7942 RNAP SI3-tail. **(B)** The structural superimposition of *Syn*7942 RNAP SI3-tail and *E. coli* SI3-NTD (PDB: 2AUK). **(C)** The protein sequence alignment of SI3-tail of *Syn*6803 and *Syn*7942 RNAP. **(D)** The protein sequence alignment of SI3-tail of *Syn*6803 RNAP and SI3-NTD of *E. coli* RNAP. The ‘RTRHG’ loop of *Syn*6803 RNAP-SI3 domain is labeled.

**Figure S5.**
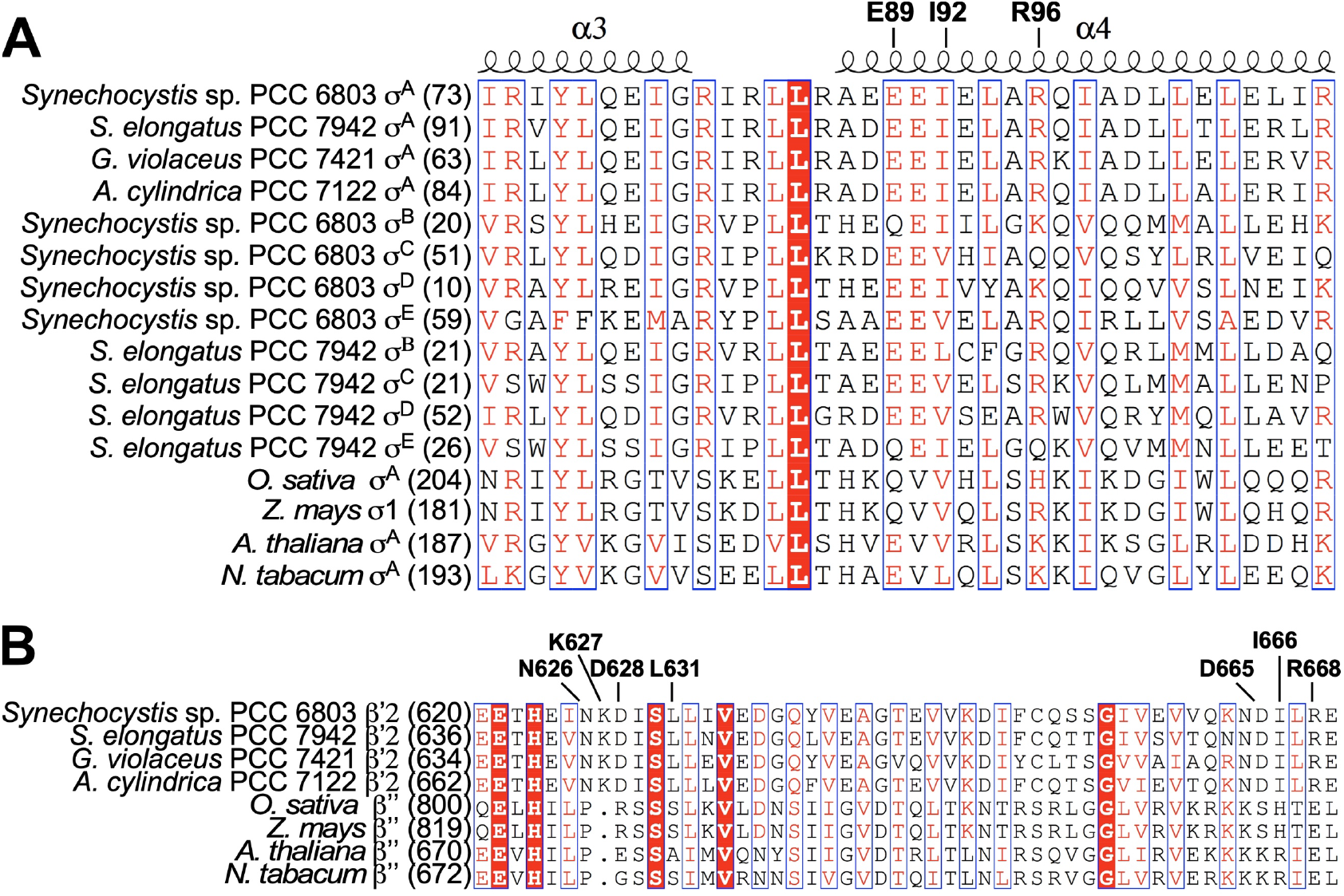
Sequence alignment of SI3-head and σ_2_. **(A)** The protein sequence alignment of the region 2 of σ factors of various cyanobacteria and plant chloroplasts. **(B)** The protein sequence alignment of RNAP SI3-head of various cyanobacteria and plastid-encoded polymerase (PEP) of plant chloroplasts.

**Figure S6.**
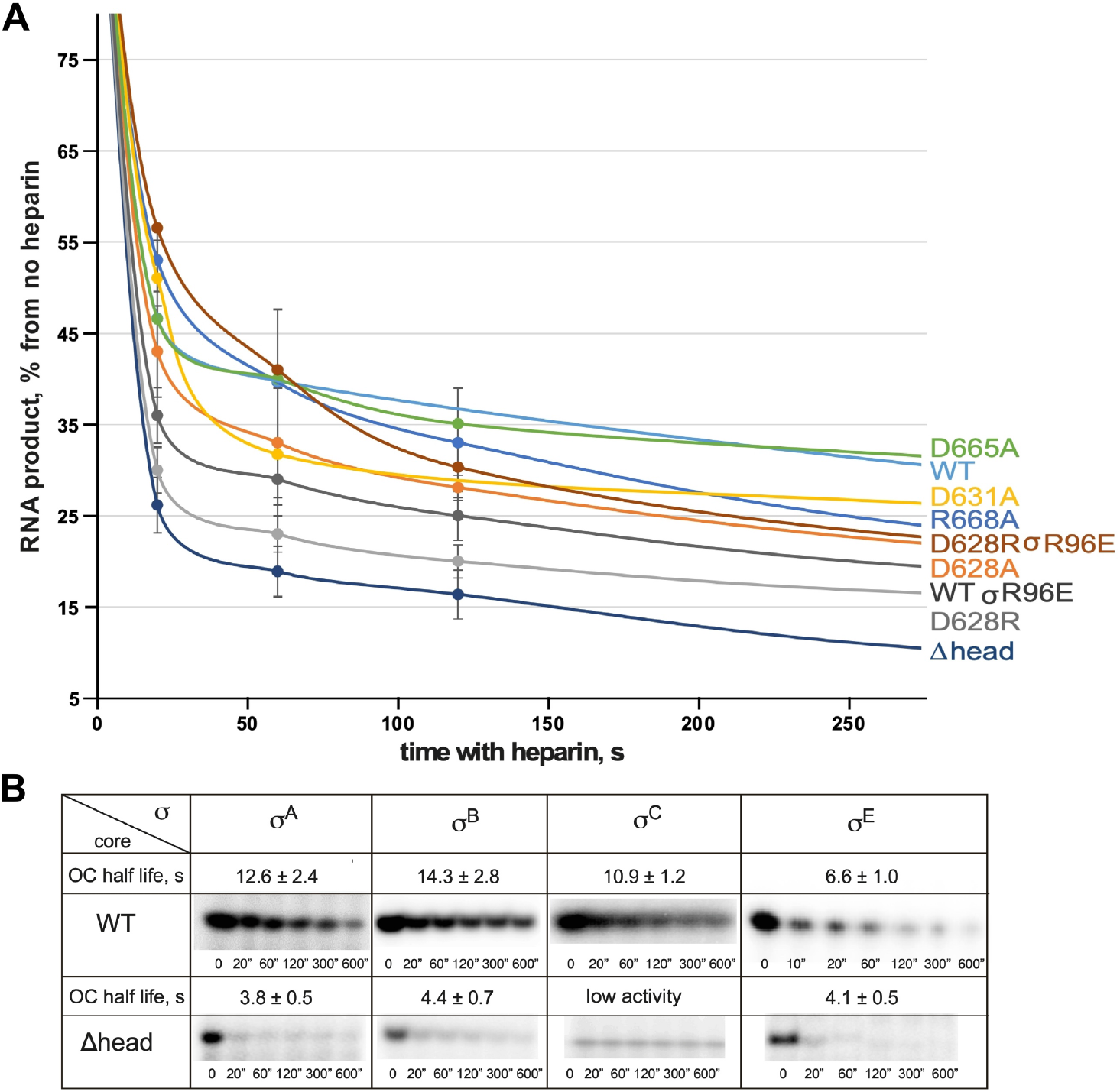
Kinetics of promoter complex decay of WT/mutant *Syn*7942 RNAP holoenzymes in the presence of heparin. **(A)** The decay cruves of WT/mutant *Syn*7942 RNAP holoenzymes in the presence of heparin. The estimated RPo half-lives is plotted in Fig. 3E. **(B)** Representative gel images and estimated *Syn*7942RPo half-lives (plotted in Fig. 3I) comprising WT/Δhead RNAP and different σ factors from three independent replicates.

**Figure S7.**
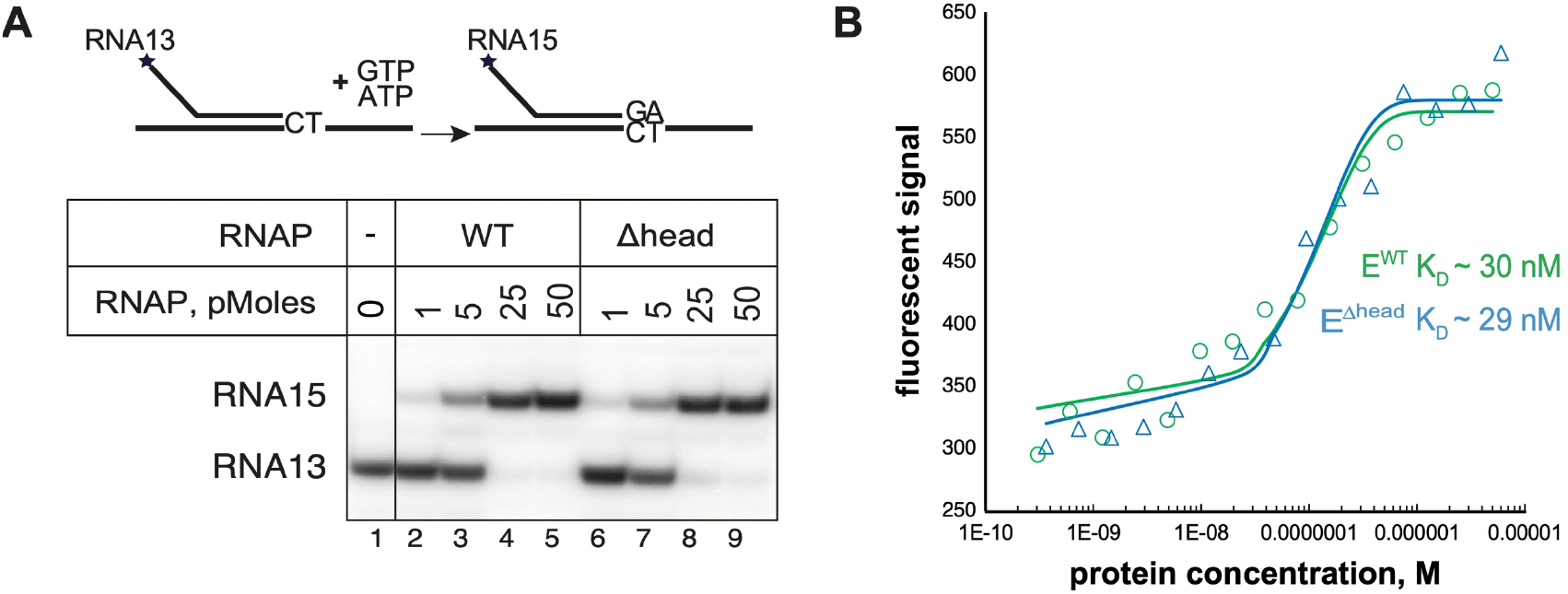
The SI3-head deletion affects neither RNAP catalytic activity nor its affinity to σ. **(A)** NTP incorporation rate is not affected upon deletion of SI3-head. The 5’ end labelled 13 nt long RNA is extended by addition of 100 µM GTP and ATP substrates to 15 nt long RNA using increased concentrations of either WT or E^βhead^ RNAP core enzymes. **(B)** Plots of concentration-dependent efficiency of holoenzyme formation and calculated dissociation constants for E^WT^ and E^βhead^ holoenzymes measured by microscale thermophoresis.

**Figure S8.**
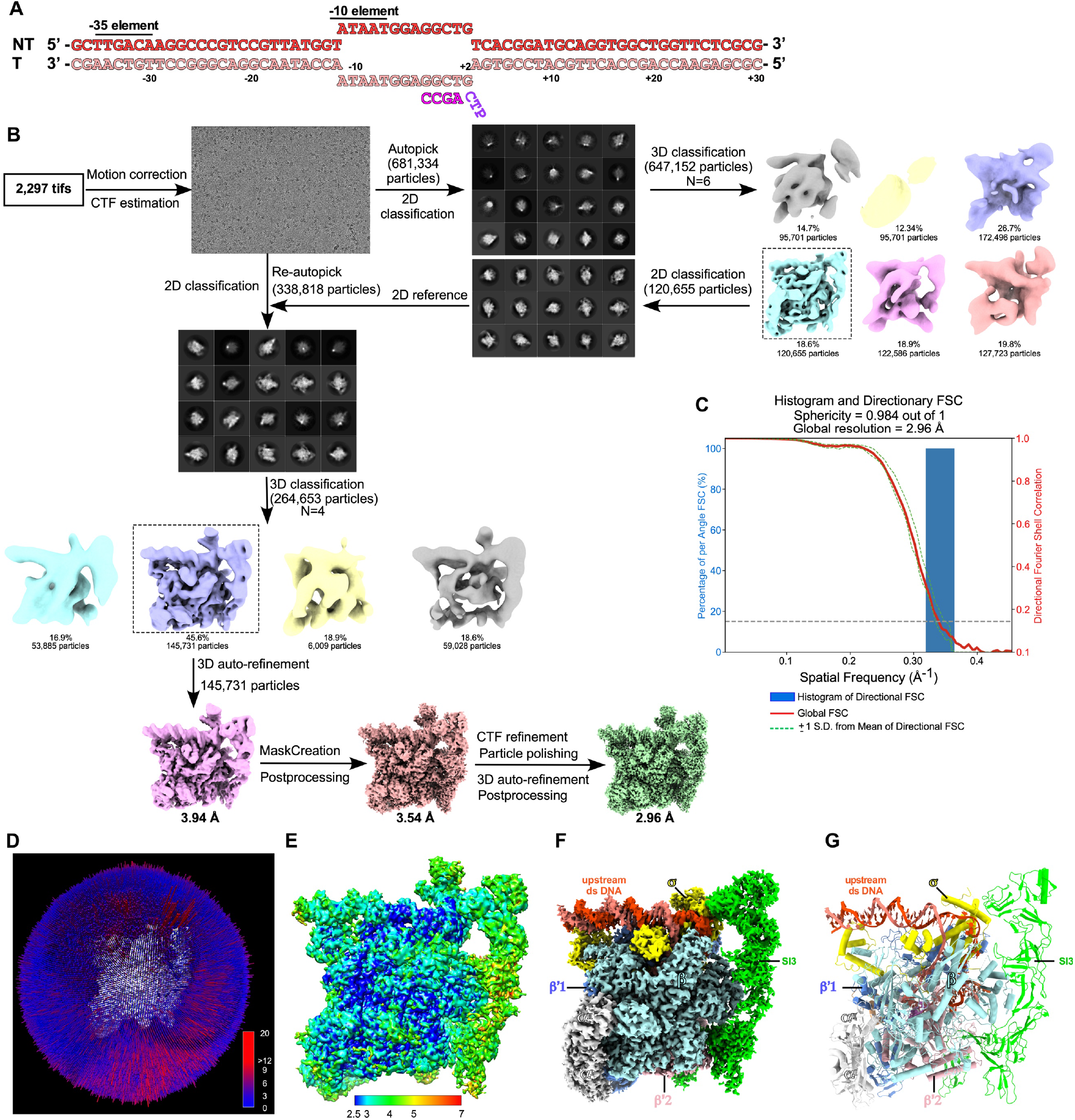
The flow chart of cryo-EM data collection and map calculation for CTP-bound RPitc. **(A)** The DNA-RNA scaffold used for cryo-EM structure determination. **(B)** The flowchart of data processing. **(C)** The 3D FSC plot. The dotted line represents 0.143 cutoff of the global FSC curve. **(D)** The angular distribution of single-particle projections by number of particles of each projection. **(E)** The cryo-EM maps of *Syn*6803 CTP-bound RPitc colored by local resolution or **(F)** by subunit. **(G)** The structure model of the CTP-bound RPitc.

**Figure S9.**
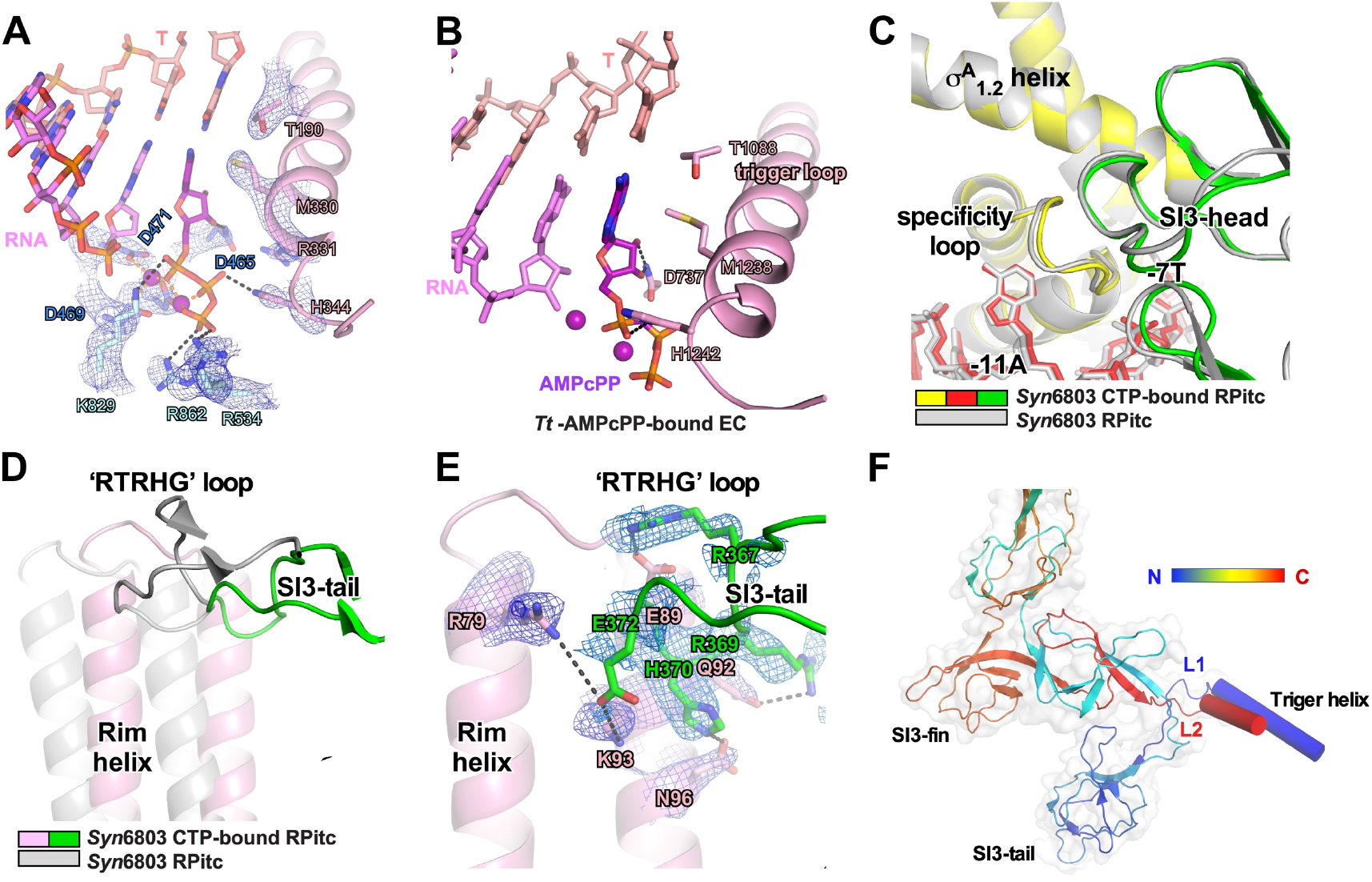
The structural analysis of CTP-bound RPitc. **(A)** The cryo-EM map and structural model of the detailed interaction between CTP and RNAP residues. **(B)** CMPcPP in the ‘i+1’ site of *Thermus thermophilus* elongation complex (PDB: 2O5J). **(C)** The structure superimposition between Syn6803 RPitc and CTP-bound RPitc shows that SI3-σ interaction remains intact upon trigger helix refolding. **(D)** The structure superimposition between Syn6803 RPitc and CTP-bound RPitc shows that the rim helices and SI3-tail move together and retain their interaction upon trigger loop folding. **(E)** The cryo-EM map of the SI3-Rim interface in the structure of CTP-bound RPitc. **(F)** The trigger loop refolding indues stretching of the two short linkers, L1 and L2 that connects the trigger helix to SI3-tail and SI3-fin domains, respectively.

**Figure S10.**
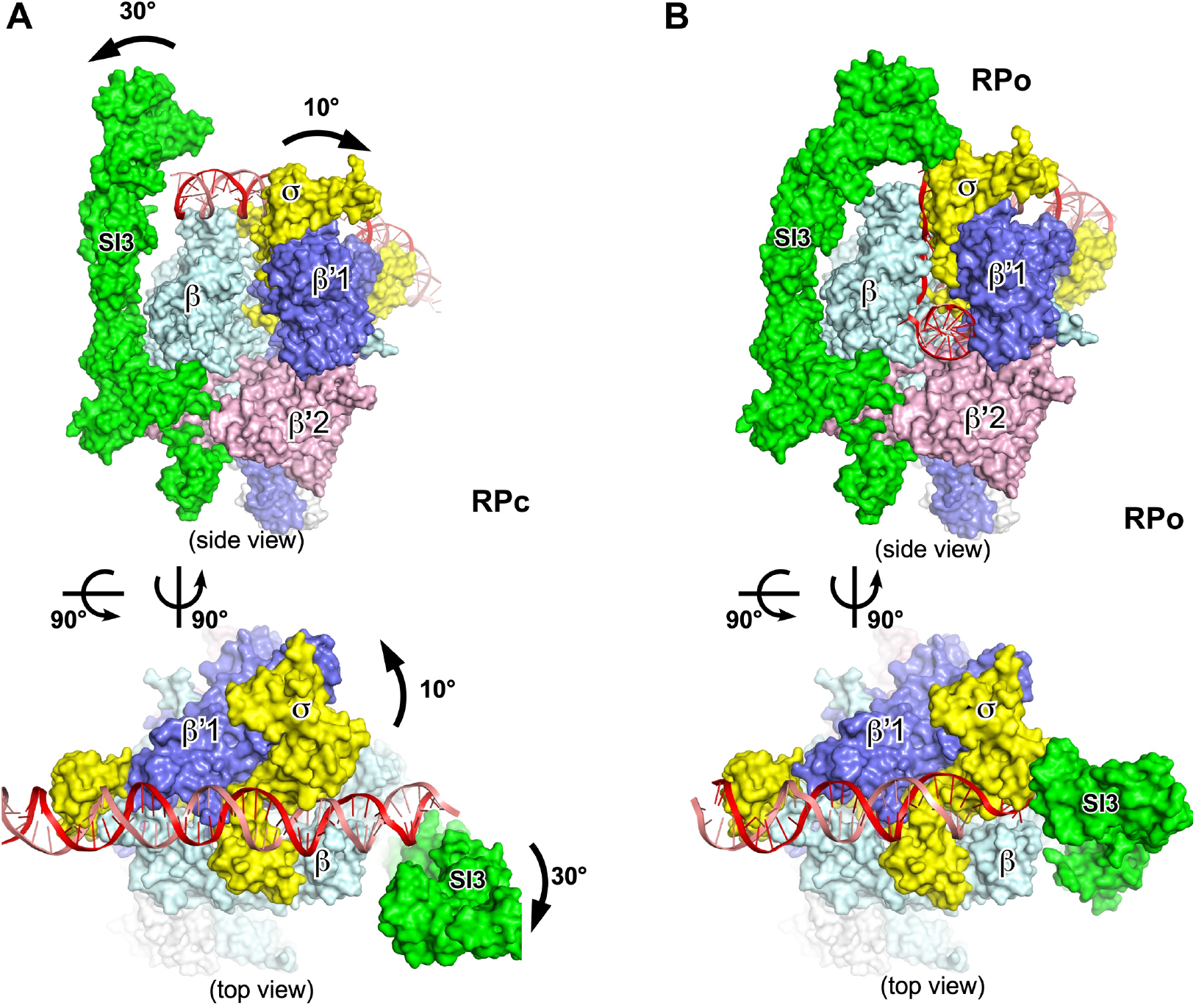
The SI3-σ arch has to open to allow loading of promoter dsDNA. **(A)** A structural model of *Syn*6803 RPc. The RNAP-SI3 and RNAP-clamp have to rotate 30° and 10°, respectively, for loading of promoter dsDNA to form a RNAP-promoter DNA closed complex (RPc). **(B)** The cryo-EM structure of *Syn*6803 RPitc, in which the SI3-σ arch is closed.

